# A model for background selection in non-equilibrium populations

**DOI:** 10.1101/2025.02.19.639084

**Authors:** Gustavo V. Barroso, Aaron P. Ragsdale

## Abstract

In many taxa, levels of genetic diversity are observed to vary along their genome. The framework of background selection models this variation in terms of linkage to constrained sites, and recent applications have been able to explain a large portion of the variation in human genomes. However, these studies have also yielded conflicting results, stemming from two key limitations. First, existing models are inaccurate in a critical region of parameter space (*N*_*e*_*s* ~ −1), where the local reduction in diversity is sharpest. Second, they assume a constant population size over time. Here, we develop predictions for diversity under background selection based on the Hill-Robertson system of two-locus statistics, which allows for population size changes. We treat the joint effect of multiple selected loci independently, but we show that interference among them is well captured through local rescaling of mutation, recombination and selection in an iterative procedure that converges quickly. We further accommodate existing background selection theory to non-equilibrium demography, bridging the gap between weak and strong selection. Simulations show that our predictions are accurate across the entire range of selection coefficients. We characterize the temporal dynamics of linked selection under population size changes and demonstrate that patterns of diversity can be misinterpreted by other models. Specifically, biases due to the incorrect assumption of equilibrium carry over to downstream inferences of the distribution of fitness effects and deleterious mutation rate. Jointly modeling demography and linked selection therefore improves our understanding of the genomic landscape of diversity, which will help refine inferences of linked selection in humans and other species.

## Introduction

Patterns of genetic variation reflect ancestral events such as historical population size changes, population splits, migrations, and episodes of natural selection. Whereas demographic processes affect the whole genome, selection operates on functionally constrained sites, sorting polymorphisms based on their phenotypic effect. The interplay between neutral stochastic processes and the deterministic force of negative selection is central to evolutionary biology (Kimura and Ohta, 1971; Cutter and Payseur, 2013; Pouyet and Gilbert, 2021; Barroso and Dutheil, 2023). Early work in population genetics theory has characterized how single-site summary statistics behave under different demographic histories and degrees of direct negative selection (Ewens, 1964; Sawyer and Hartl, 1992; Marth et al., 2011).

The effects of negative selection extend beyond constrained sites. This is because physical linkage connects the evolutionary histories of sites along the same chromosome, such that linked loci tend to share coalescence events en route to their most recent common ancestors (Hudson and Kaplan, 1988, 1995b). On the other hand, recombination weakens these correlations by separating lines of descent at crossing-over breakpoints during meiosis. Consequently, since the rate of recombination between loci increases with their physical distance, nearby sites share a larger fraction of their histories than sites that are farther apart (Hill and Robertson, 1966, 1968; Ohta and Kimura, 1971; Cutter and Payseur, 2013). The distortion in genealogies of neutral sites caused by the presence of negatively constrained loci is termed background selection and has received much attention in the past three decades (e.g., Charlesworth et al., 1993; Hudson and Kaplan, 1995b; Nordborg et al., 1996; McVicker et al., 2009; Good et al., 2014; Cvijović et al., 2018; Elyashiv et al., 2016; Ewing and Jensen, 2016; Torres et al., 2020; Murphy et al., 2022; Buffalo and Kern, 2024). In genomic data, a well-documented signature of background selection is the spatial covariance between genetic diversity and features like gene density and recombination rate (Aguade et al., 1989; Begun and Aquadro, 1992; Andolfatto and Przeworski, 2001; Lohmueller et al., 2011; Castellano et al., 2018). Accurately characterizing the evolutionary mechanisms that shape these patterns is key to establishing a proper baseline model of genome evolution (Comeron, 2017; Johri et al., 2020).

The most well-studied measure of background selection is the reduction in pairwise nucleotide diversity, *π*. Classic background selection theory (cBGS) describes how the ratio *π/π*_0_ (denoted *B*, with *π*_0_ being the expected diversity in the absence of linked selection) is affected by the continual influx and removal of strongly deleterious variants (Charlesworth et al., 1993; Hudson and Kaplan, 1995a; McVicker et al., 2009). The assumption of mutation-selection balance employed by cBGS offers elegant formulas which incorporate multiple constrained sites through simple multiplication. At the time, the approximations of cBGS were justified by the prevailing view that the strength of selection in nature should be quite high, with selection coefficients against heterozygous genotypes on the order of −0.01 and stronger. Under this regime, selection is strong enough that its effect can indeed be treated deterministically (Haldane, 1927), even in relatively small populations, and equilibrium diversity at constrained sites is low enough that they do not substantially interfere with each other, even at short genetic distances (Hill and Robertson, 1966). Thirty years on, however, investigation of genomic data from several species has revealed wide distributions of fitness effects, with mean selection coefficients on the order of −0.001 and typically long tails spanning from moderately selected to nearly-neutral mutations (Eyre-Walker et al., 2006; Eyre-Walker and Keightley, 2007; Huber et al., 2017). Furthermore, other empirical work has found evidence of pervasive interference among selected sites (Castellano et al., 2016).

Yet predictions of *B*-value maps (*B*-maps) in humans and fruit flies have relied on the analytical approximations of cBGS (McVicker et al., 2009; Elyashiv et al., 2016; Murphy et al., 2022). Only recently has a different theoretical framework (Santiago and Caballero, 1995, 1998, 2016) been translated into the computational machinery required for statistical inference (Buffalo and Kern, 2024). Unlike cBGS, the approach from Santiago and Caballero (2016) (SC16) is a quantitative model based on the variance in fitness within a population, which is a function of the influx of deleterious variation and rate of fixation of slightly deleterious mutations (i.e., Muller’s Ratchet, Muller, 1964). In practice, this has produced accurate predictions of *B*-values in the weak selection regime (Buffalo and Kern, 2024). However, the model (as currently implemented) breaks down for scaled selection coefficients (*N*_*e*_*s*) in the broad vicinity of −1. This is a particularly compelling region of parameter space, where interference among constrained sites is highest and the local reduction in diversity is strongest (McVean and Charlesworth, 2000; Comeron and Kreitman, 2002; Good et al., 2014; Ragsdale, 2022). More realistic models of linked selection rely on accurate prediction of *B*-values in this regime.

Current models have other limitations besides the range of selection regimes where they apply. A critical shortcoming shared by cBGS and SC16 theory is their restriction to equilibrium demography. Fluctuating population sizes impact the segregation trajectories of deleterious variants, which in turn influence diversity at linked neutral sites. Predicting these dynamics is challenging because the drift-effective population size (*N*_*e*_) simultaneously dictates the overall level of (deleterious) genetic variation maintained in the population (*O*(*N*_*e*_*µ*)), the relative strength of selection versus drift (*O*(*N*_*e*_*s*)), and the time available for recombination to break down correlations among genealogies (*O*(*N*_*e*_*r*)). For example, deleterious variants subject to efficient selection in a large ancestral population may transition to the “interference” regime (*N*_*e*_*s* ~ −1) after a sharp bottleneck, where drift plays a pivotal role. Meanwhile, concerted decreases in *N*_*e*_*µ* and *N*_*e*_*r* will lead to lower deleterious diversity and tighter linkage, respectively, with opposing effects on the extent of linked selection. What, then, is the net outcome of these perturbed forces on *B*-values?

The distinct relationships of *B* against *N*_*e*_*s, N*_*e*_*r* or *N*_*e*_*µ* would be enough to challenge intuition, but the situation is further complicated because the drift-effective population size, as recorded in genetic diversity, represents the harmonic mean between the sizes in consecutive epochs (Charlesworth, 2009). Therefore, after a sudden demographic shift and until the new equilibrium is reached, *N*_*e*_ itself changes gradually over time, reflecting the proportion of uncoalesced ancestry within each epoch, and with it the compound parameters *N*_*e*_*s, N*_*e*_*r* and *N*_*e*_*µ*. In general, if the time-scale of consecutive demographic changes is short enough to prevent the population from reaching stationarity, as is the case in natural populations, then patterns of diversity observed at any time should arise from a complex combination of past dynamics and not simply their average. While previous work used non-equilibrium simulations to describe the temporal dynamics of *B*-values (Torres et al., 2020), incorporating them into theoretical models is important for the development of inference tools that can be applied to data.

In this article we introduce a new model (`moments++`, Barroso and Ragsdale, 2025) that can accurately predict *B*-values under a wide range of evolutionary scenarios, from weak to strong selection, passing through the interference selection regime, and under non-equilibrium single-population demography. Our method is based on the Hill and Robertson (1968) system of two-locus statistics that has recently been generalized (Ragsdale and Gravel, 2019). We incorporate selection into the two-locus system and model its effect on a neutral locus located at an arbitrary recombination distance. Inspired by the pioneering work of Hudson and Kaplan (1995a,b), we treat the joint effect of multiple selected loci independently, but we show that under equilibrium, interference among them is well captured through iterative rescaling of mutation, recombination and selection. We explore the dynamics of linked selection under non-equilibrium demography and discuss how patterns of diversity can be misinterpreted by other models.

## Results and Discussion

### Predicting diversity under linked selection in a two-locus framework

Our objective is to model the effect of linked selection on the expected diversity of a focal site. We begin our approach with a pure two-locus model, where a single constrained site stands a recombination distance *r* away from the focal neutral site (Appendix A, Figure 1A). Although *B*-values are agnostic to possible selection on the focal site, we develop the two-locus model under the assumption that the focal site evolves neutrally to avoid formalizing its reciprocal linked selection effect. Later, using a heuristic correction for multi-locus interference, we will relax this assumption and predict diversity for neutral and constrained sites across the genome.

**Figure 1.**
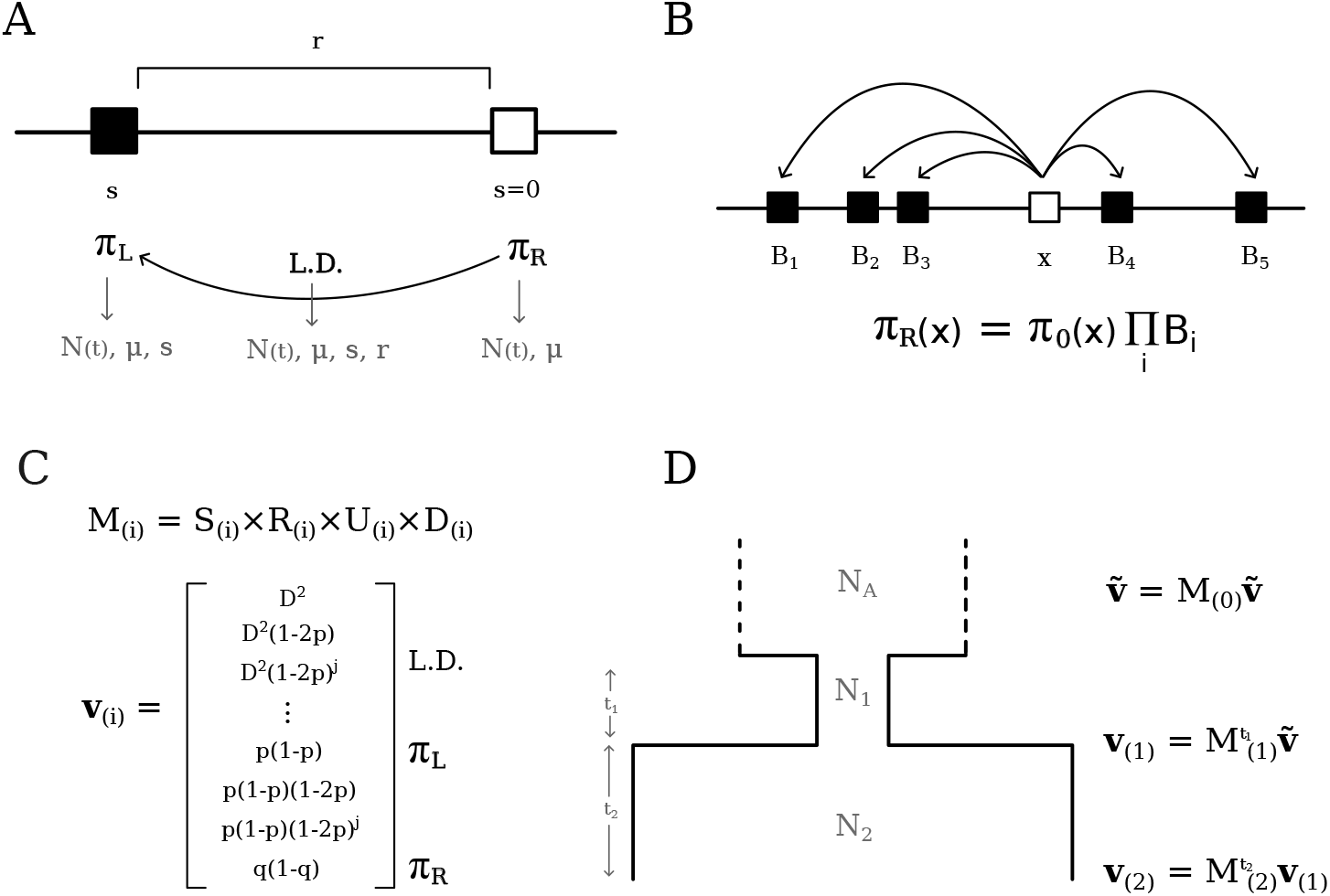
Schematic representation of the Hill-Robertson system with negative selection against the derived allele at the left locus. A) In a pure two-locus model, neutral pairwise diversity (*π*_*R*_) is modulated by LD with a single constrained site located at recombination distance *r* that experiences selection strength *s*. B) The multi-locus extension of the pure two-locus model, where the total reduction in diversity, arising from many pairwise interactions, is multiplicative. C) A simplified representation of the system components, where *M* is the transition matrix that embodies the evolutionary parameters and operates on **v** (*i* indexes the epoch). D) Obtaining expectations under a three-epoch model, starting from equilibrium in the ancestral population (top) and finishing at present time (*t*_*i*_ is the number of generations spent on epoch *i*).

Negative selection in the constrained locus interferes with the expected sojourn time at the focal neutral locus. Briefly, the presence of a deleterious variant introduces fitness variance in the population, such that neutral polymorphisms associated with the deleterious background have increased extinction probability whereas those associated with the ancestral background have increased chance of fixation. The magnitude of this effect depends on the extent of linkage disequilibrium (LD) between the constrained and neutral loci, suggesting the two-locus model of Hill and Robertson (1968) as a natural framework for our endeavor.

In the absence of linked selection, expected pairwise diversity at a neutral locus, E[*π*_0_], is governed by a balance between mutation and drift (Wright, 1931). This follows the familiar expressions for the expected changes in diversity in one generation:

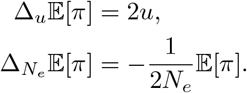

The dynamics of 𝔼 [*π*] under neutrality depend only on itself, and this simple system closes. With selection at a linked locus (left, arbitrarily), allele frequencies and pairwise diversity at the neutral (right) locus change depending on its association with the selected allele (Appendix B). For example, the expected allele frequency (*q*) at the right locus changes due to selection at the left locus as Δ_*s*_ 𝔼 [*q*] = *s* 𝔼 [*D*], where *D* is the standard covariance measure of LD. For pairwise diversity at that locus,

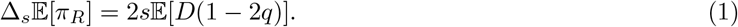

Here, 𝔼 [*D*(1 − 2*q*)] is positive, and the decay of *π*_*R*_ due to LD with the constrained locus becomes explicit.

With the inclusion of linked selection, the number of statistics needed to model 𝔼 [*π*_*R*_] grows and the system and no longer closes (Appendix B). This happens because the addition of 𝔼 [*D*(1 − 2*q*)] also requires understanding its dynamics under mutation, drift, recombination and selection (Ragsdale and Gravel, 2019). In computing these terms, the system of recursions on these statistics (moments of the full two-locus haplotype frequency distribution) accumulates an ever-increasing number of moments, with the presence of selection consistently demanding moments of slightly higher order than those previously found (Appendix A, and see Jouganous et al. (2017); Ragsdale and Gravel (2019)). However, these take a familiar form, as we soon find moments that were originally studied by Hill and Robertson (1968); Ohta and Kimura (1969) to compute the variance of *D* under neutrality: 𝔼[*D*^2^], 𝔼[*D*(1 − 2*p*)(1 − 2*q*)], and 𝔼[*p*(1 − *p*)*q*(1 − *q*)] (where *p* and *q* are the derived allele frequencies at the left and right loci, respectively). Therefore, the moments accumulated due to the inclusion of selection resemble those in the original Hill-Robertson basis, augmented by factors of (1 − 2*p*)^*j*^, with *j* ≥ 0. That is, the system includes all terms 𝔼[*D*^2^(1 − 2*p*)^*j*^], 𝔼[*D*(1 − 2*p*)^*j*^(1 − 2*q*)], and 𝔼[*p*(1 − *p*)(1 − 2*p*)^*j*^*q*(1 − *q*)], as well as the single-locus terms 𝔼[*π*_*L*_(1 − 2*p*)^*j*^], and we label this basis **v**. Within **v**, each moment with a given factor (1 − 2*p*)^*j*^ depends only on its close neighbors (moments with 1 − 2*p* factors of *j* ± 1 or 2) so that the system remains sparse. This suggests that a treatment of the two-locus system with selection is manageable, but a moment-closure approximation is required to obtain a concise matrix representation of the recursions.

Surprisingly, naive truncation of the system after a sufficient number of 1 − 2*p* factors produces accurate results (Figure S7). Since stronger selection demands faster replacement of eliminated haplotypes (Appendix A, the required maximum order of (1 − 2*p*)^*j*^ is an increasing function of |*N*_*e*_*s*|. Since the extra information brought by the inclusion of a new haplotype in a small sample is higher than in a sample that is already large, this function is supra-linear (Jouganous et al., 2017). Throughout the remainder of this article, when modeling different strengths of selection, we compute statistics using the appropriate truncation level of (1 − 2*p*)^*j*^ factors needed for accurate prediction, but omit this information for ease of exposition.

### Modeling strong linked selection under non-equilibrium demography

The demanding nature of the selection operator imposes practical problems for |*s*| ≳ 𝒪 (0.01) (see Conclusions). Fortunately, the strong-selection regime is where cBGS approximations can take over without losing accuracy (Charlesworth et al., 1993; Hudson and Kaplan, 1995b; Nordborg et al., 1996). In particular, Nordborg (1997) showed that the structured coalescent framework can be used to model background selection through a separation of time scales. In this approach, coalescence at the neutral locus occurs within allelic classes defined by the number of haplotypes in a sample that carry the deleterious variant at the constrained locus, and transitions between classes due to recombination and mutation are rapid relative to coalescence. With strong selection, mutation-selection balance applies, maintaining the frequency of the deleterious class at *p* = *u/s*, and the reduction in coalescence times was found to be independent of *N*_*e*_ (Nordborg et al., 1996; Nordborg, 1997):

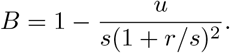

To incorporate population size changes, we assume a piecewise-constant population size history. Using phase-type theory (Hobolth et al., 2019), we can efficiently calculate probabilities of coalescence within each epoch and the expected *T*_*MRCA*_ conditional on coalescing within a given epoch (Methods). This is done for both the standard neutral coalescent and structured coalescent (Nordborg, 1997) models to obtain the expected reduction in diversity.

On the surface, the independence of *B*-values from *N*_*e*_ at steady state could suggest that they would be invariant to changes in population size. Indeed, size changes have little effect on deleterious diversity, so that *π*_*L*_ remains approximately constant (Figure S8, top panels). However, *B*-values are transiently affected, as both *π*_0_ and *π*_*R*_ depend more sensitively on *N*_*e*_ – and to different degrees. To see this, consider the distribution of genealogies under a two-epoch demographic model. After the size change, and until steady state is re-established, the probability of coalescence within the most recent epoch disproportionally differs whether the neutral locus is linked to a selected locus or not. This distorts the reduction in the expected coalescence times of linked neutral sites relative to unlinked sites, so that the ratio *π*_*R*_*/π*_0_ varies temporally. Another interpretation of this phenomenon is that *π*_*R*_ reaches the new steady-state at a faster rate than *π*_0_ due to its increased coalescence rate, since to a first approximation, linked selection incurs a reduction in *N*_*e*_ (Charlesworth et al., 1993; Charlesworth, 2009). Figure S3 demonstrates this effect, and our theoretical results confirm observations from simulations carried out in Torres et al. (2020).

The extension of the Nordborg (1997) cBGS framework to non-equilibrium demography completes the bridge between weak and strong selection. By joining the numerical solutions of moments++ and the analytical predictions extended from Nordborg’s model, we can predict *B*-values under arbitrary single-population demographic histories and distributions of fitness effects. While we focus here on changes in population size, this approach based on phase-type theory is quite flexible. It may be extended to accommodate piecewise-constant changes in the mutation rate at the selected site, recombination rate, the strength of selection (as long as it remains strong, relative to drift), or a combination of these.

### *B*-value behavior under equilibrium demography

We first benchmarked our model against equilibrium (*N*_*e*_ = 10, 000) simulations of a 100 kb segment with uniform recombination and mutation rates (Methods). In multi-locus settings, the *B*-value of each site is obtained by multiplying together the *B*-values resulting from every pairwise interaction. Therefore, our approach follows the formulation of Hudson and Kaplan (1995a,b), who started from two-locus reasoning to arrive at analytical approximations of multi-locus effects. We observe close agreement between our predictions and simulations, from weak to strong selection, with substantial overlap in the regime where cBGS can take over (Figure 2). Moreover, unlike the implementation of Buffalo and Kern (2024), there is no discontinuity in the prediction around *N*_*e*_*s* of −1. This is precisely the peak of linked selection, where the combination of intermediate deleterious diversity and intermediate rates of selective elimination leads to a sharp reduction in diversity at nearby linked sites.

**Figure 2.**
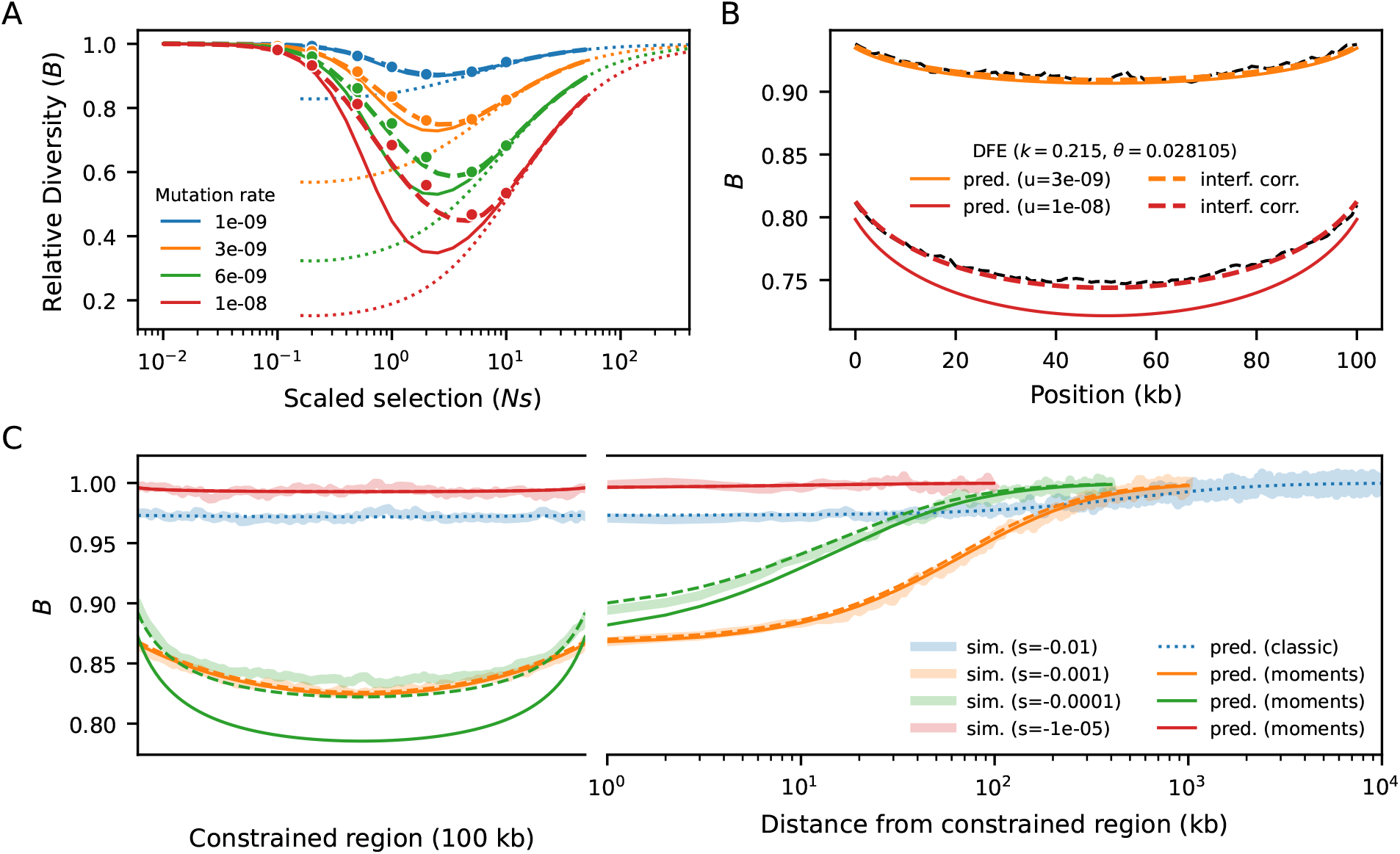
Benchmarking against equilibrium simulations of a 100 kb segment (*N*_*e*_ = 10, 000). A) *B*-value in the middle of the segment, for different mutation rates and strengths of selection. Dots are simulated *B*-values (averaged over 10,000 replicates). Solid lines show naive predictions, dashed lines incorporate interference correction and dotted lines mark cBGS. B) *B*-map predictions remain accurate when *s* follows a distribution of fitness effects (black dashed lines show simulated data). C) *B*-values along the simulated segment. A concave pattern emerges because loci near the edges are on average further from constrained elements, and is counteracted by stronger interference in the center, with flattens the *B*-map. As the distance from the selected segment increases moving into a neutral flanking region, linked selection relaxes at a pace that depends on *s* and *r*. Weak-to-moderate selection (*N*_*e*_*s* ~ −1) distorts patterns of diversity sharply albeit locally, whereas stronger selection (*N*_*e*_*s* ≤ −10) imposes a small but still relevant reduction throughout the entire region. Solid lines show naive predictions, dashed lines incorporate interference correction, and shaded envelops mark simulated data.

Because our multiplicative model assumes that constrained loci evolve independently, ignoring interference leads to artificially deflated predictions at higher mutation rates. To alleviate this bias, we draw inspiration from cBGS (Charlesworth et al., 1993; Charlesworth, 2009) and model interference as a reduction in *N*_*e*_ at constrained loci, which varies along the genome. In practice, we simultaneously scale *µ, r* and *s* by the corresponding *B*-values at each locus (Good et al., 2014), such that the effective rates of mutation, recombination and the efficiency of selection reflect local *N*_*e*_ across the genome. This changes *B*-value predictions all around, so we repeat the process until convergence (Methods, Figure S1). This procedure restores accuracy, and we conclude that `moments++` can predict *B*-values over a wider range of selection strengths than previous methods.

We now increase the complexity of the simulations by partially mimicking the first 30 Mb of human chromosome 2 (Methods). The heterogeneous distributions of constrained elements (here, annotated exons (Schneider et al., 2017)) and recombination rates (Spence and Song, 2019) lead to intricate patterns of linked selection along this segment, which our model predicts remarkably well (Figure 3). Regions that are on average distant from exons experience weaker reduction in diversity and negligible interference. However, even in regions largely devoid of exons, *B*-values stand unequivocally below one. This happens because they are affected by strongly selected sites whose effect is less sensitive to genetic distance (Figure 2C). Conversely, *B*-values dip in exon-rich regions, especially if the local recombination rate is low. This is driven by the strong narrow-range distortions caused by weak-to-moderate selection (*N*_*e*_*s* ~ −1). These results suggest that predicting *B*-values across the full range of selection coefficients – rather than truncating the distribution of fitness effects (DFE) to exclusively consider strong selection (McVicker et al., 2009; Murphy et al., 2022) – can substantially improve models of genetic diversity (Figure S6).

**Figure 3.**
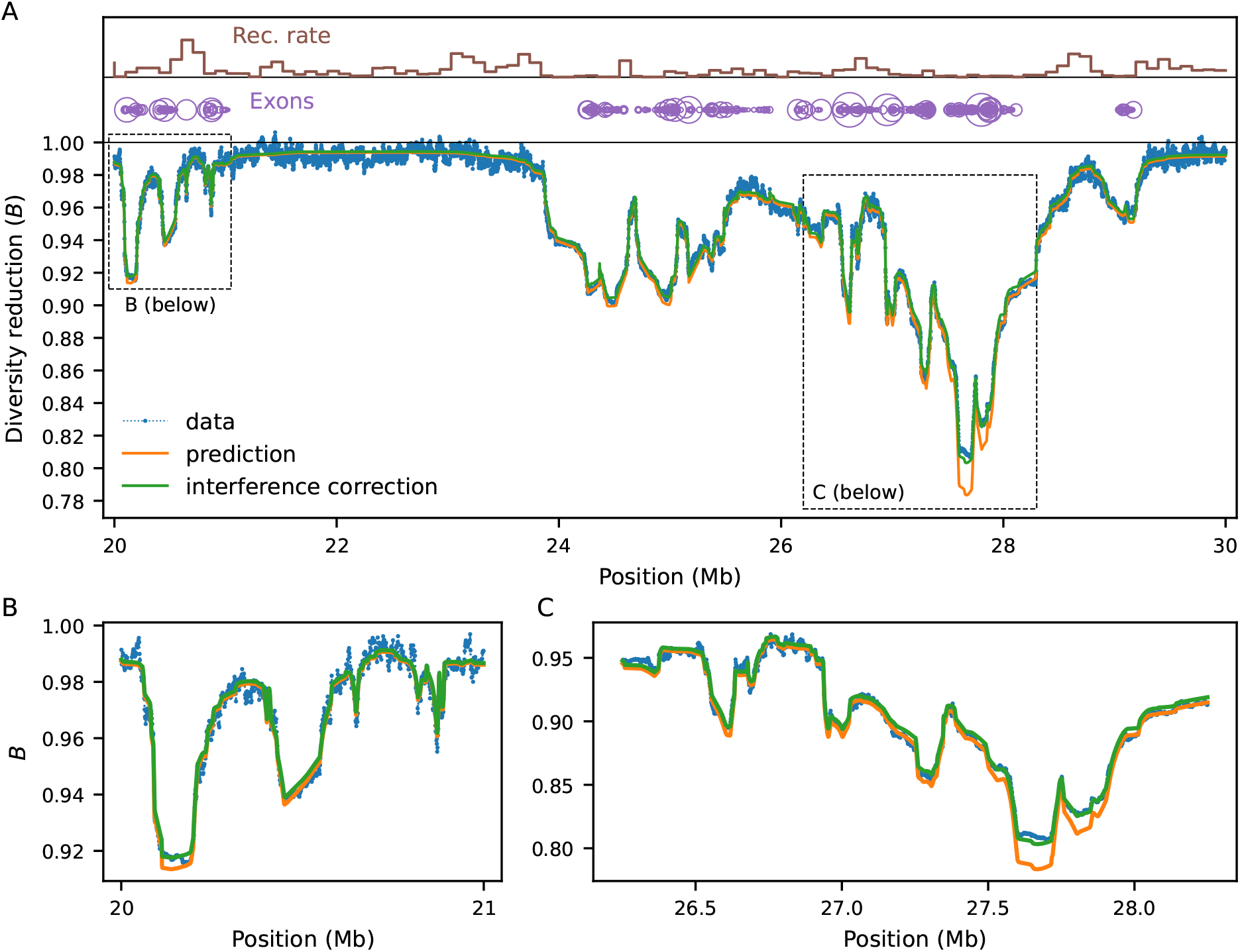
Benchmarking against equilibrium simulations (*N*_*e*_ = 10, 000) mimicking the structure of human chromosome 2, and zooming in on a 10 Mb segment (A). Local recombination rates and exon density are shown at the top. Simulated data (averaged over 10,000 replicates) is shown by blue dots. Green and orange lines show prediction with and without interference correction, respectively. Regions distant from exons experience weaker reduction in diversity, whereas *B*-values dip in exon-rich regions, especially if the local recombination rate is low (cf. panels B and C). Here the mutation rate is 10^−8^ and *s* follows the Kim et al. (2017) DFE.

### *B*-value dynamics in non-equilibrium populations

At steady-state, genetic diversity is balanced among drift, mutation, recombination and (linked) selection. Changes in population size lead to transient dynamics of linked selection, until the new equilibrium is reached (Torres et al., 2020). Since *B*-values represent the accumulated reduction in diversity at a focal site due to 𝒪 (*N*_*e*_) generations of negative selection on linked sites, and since *N* (*t*) concertedly modulates the efficiency of selection, deleterious diversity and effective recombination, we find that population fluctuations can have a substantial impact on the evolution of this statistic. *B*-value dynamics are often non-monotonic over time, depending on the strength of selection and demographic history (Figures S9, S10, Appendix C).

Here we focus on multi-locus models, in which we couple the chromosome layout from Figure 3 (including a DFE inferred from human data by Kim et al. (2017)) with three demographic models: 1) a 10-fold bottleneck (*N* = 10 000 to 1 000, tracked for 25,000 generations after size change); 2) a 10-fold expansion (*N* = 10 000 to 100 000, also for 25,000 generations); and 3) a four-epoch model that loosely resembles human effective population size history inferred by Cousins et al. (2024). By sampling the expected *B*-map at different time points after the first population size change, we show that neglecting demography can bias predictions and subsequent downstream inferences.

To assess the bias introduced by incorrectly assuming equilibrium, we used the drift-effective population size at each sampling generation (*N*_*e*_(*t*) = *π*_0_(*t*)*/*4*µ*) to predict a series of steady-state *B*-maps. This approximation assumes that deleterious diversity and selective efficiency have remained constant over time, as well as the population-size scaled recombination and mutation rates. Therefore, they contrast with the demography-aware *B*-map which reflects genealogical history and evolves over time.

In the bottleneck scenario, the discrepancy between equilibrium and demography-aware *B*-maps initially grows as the ancestral size contributes less to drift-effective *N*_*e*_ and then shrinks again as *N*_*e*_ approaches the current size (Figure S4, blue). The bias peaks ~ 5, 000 generations after the bottleneck, when most of the change in *π*_0_ has already occurred (drift-effective *N*_*e*_ ~ 2, 000), suggesting an asymmetric effect of historical population sizes in shaping *B*-values. Conversely, the expansion scenario sees the bias grow throughout the 25,000 generations of evolution (Figure S4, orange), whereas the Cousins et al. (2024) model shows more subtle deviations from equilibrium *B*-maps (Figure S5). Since in the latter model the ratios between consecutive population sizes do not differ as dramatically as a 10-fold bottleneck or expansion, discrepancies in predicted *B*-maps are not as pronounced. Thus, assuming equilibrium will bias predictions in general, in agreement with simulations (Torres et al., 2020). However, the magnitude of the bias depends on the precise population history and DFE, such that equilibrium *B*-maps may be a reasonable approximation in some systems.

### Implications for DFE inference

What are the practical consequences of neglecting demography when predicting *B*-values? It seems reasonable that the ensuing biases should carry over to downstream analyses. In this regard, previous applications of equilibrium models of linked selection have also estimated selection parameters, including the DFE and deleterious mutation rate, by minimizing the distance between the predicted landscape of diversity and that observed in human data (McVicker et al., 2009; Murphy et al., 2022; Buffalo and Kern, 2024).

These estimates have been subject to debate. Specifically, the inferred (deleterious) mutation rate is much larger than expected, incompatible with more direct estimates of the human mutation rate (Ségurel et al., 2014; Tian et al., 2022)). Furthermore, the DFE of functional elements such as exons conflicts with those obtained from the site frequency spectrum (Williamson et al., 2005; Keightley and Eyre-Walker, 2007; Boyko et al., 2008; Kim et al., 2017). For some classes of constrained elements the inferred DFE is strongly bimodal, implying a combination of linked selection that is either weak and very localized (|*N*_*e*_*s*| ≪ 1) or long range (|*N*_*e*_*s*| ≫ 1). This suggests problems with identifiability: the elevated mutation rate increases deleterious diversity, reducing *B* and partially compensating for the low density of mutations around *N*_*e*_*s* = −1.

To investigate the bias introduced by assuming equilibrium, we examined scenarios of fluctuating demography and several DFE parameterizations. We predicted *B*-maps in the ancestral population and at every 500 generations after the first population size change, which we then used to infer the deleterious mutation rate and the shape and scale of the Gamma-distributed DFEs that best fit those maps under the assumption of steady-state size history. At each sampling time point, the maximum-likelihood estimates of the aforementioned selection parameters will be those which minimize the distance between equilibrium and demography-aware *B*-maps. This setup provides a best-case scenario for parameter inference by removing any other source of error, technical or biological (including genealogical noise), that would be present in an analysis of real data, and in this sense it offers a conservative assessment of the bias.

Figure 4 compares simulated and inferred parameter values for the two-epoch scenarios. As expected, they are well recovered in the ancestral population which finds itself at steady-state. As time moves forward after the size change, a strong departure develops under both bottleneck and expansion. The direction of the bias is opposite between demographic scenarios, and within each it is again opposite between shape and scale of the DFE. Mirroring the patterns observed in the previous subsection, the bias grows as the ancestral size contributes less to drift-effective *N*_*e*_. Under the bottleneck, it then shrinks as *N*_*e*_ approaches the current size (Figure 4, blue curves). Conversely, in case of expansion, it grows throughout the 25,000 generations observed, and predictions from the pure two-locus model (Appendix C, Figure S10) suggest that it would take a long time to restore accuracy using equilibrium assumptions. We also see that the magnitude of the bias depends on the selection regime, with the shape being the most sensitive parameter. Translating inferences to the mean and variance of the DFE, stronger selection leads to stronger absolute deviations (Figure S11).

**Figure 4.**
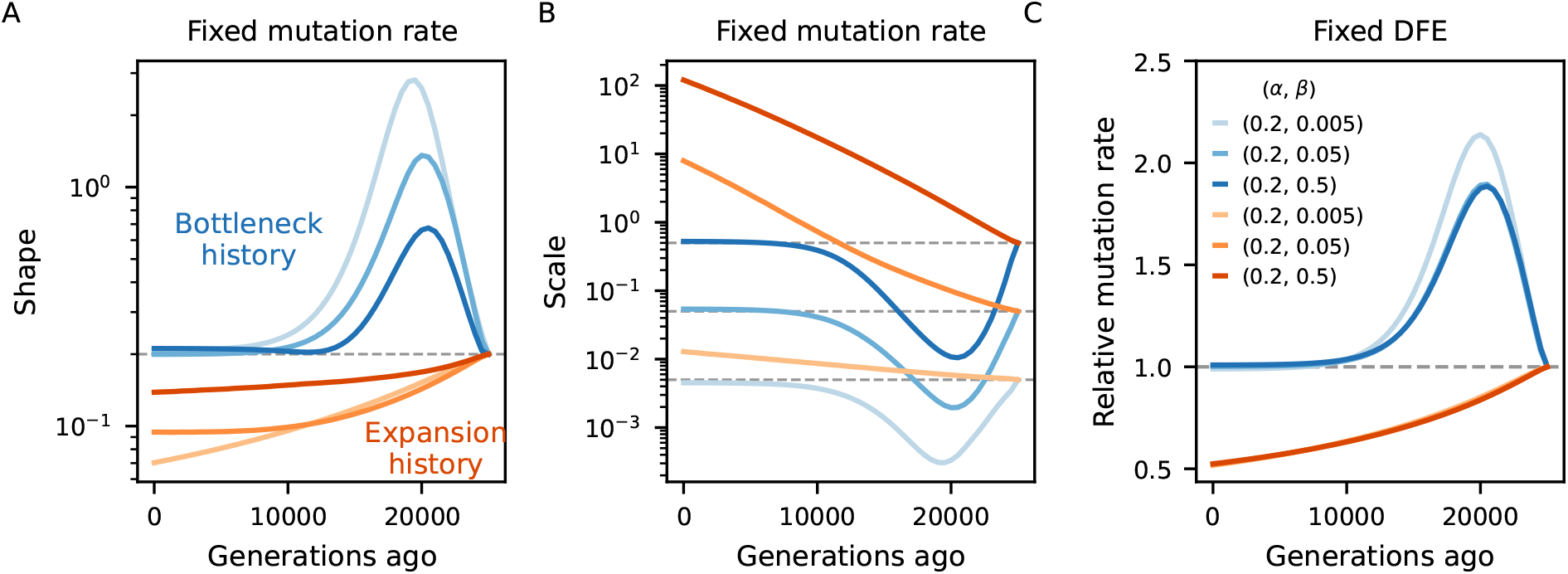
Biases in the inference of selection parameters due to the incorrect assumption of equilibrium. An ancestral population experiences either a 10-fold bottleneck (Blues) or expansion (Reds) and is followed for 25,000 generations after the size change. Solid lines in A-B show the inferred shape and scale of the DFE at different points in time (*N*_*e*_(*t*) = *π*_0_(*t*)*/*4*µ*), which flows from right to left (dashed lines denote simulated value). Solid lines in C show the relative inferred (deleterious) mutation rate. We either assume the mutation rate is known and fit the shape and scale of the DFE (A-B), or vice-versa (C).

The behavior is markedly different in the human-like history (Cousins et al., 2024), where consecutive population sizes move the target equilibrium point in opposite directions. Here we tested more dissimilar DFEs and found that the shape parameter is underestimated under strong selection but overestimated otherwise (Figure 5B). The mean selection coefficient is underestimated, but the precise temporal trajectory of the bias depends on the DFE (Figure S12). Since the out-of-Africa event is relatively recent (Gutenkunst et al., 2009; Gravel et al., 2011), we expect that *B*-maps from distinct human populations will be fairly similar, even after accounting for demography, in agreement with what has been found under equilibrium (Murphy et al., 2022; Buffalo and Kern, 2024).

**Figure 5.**
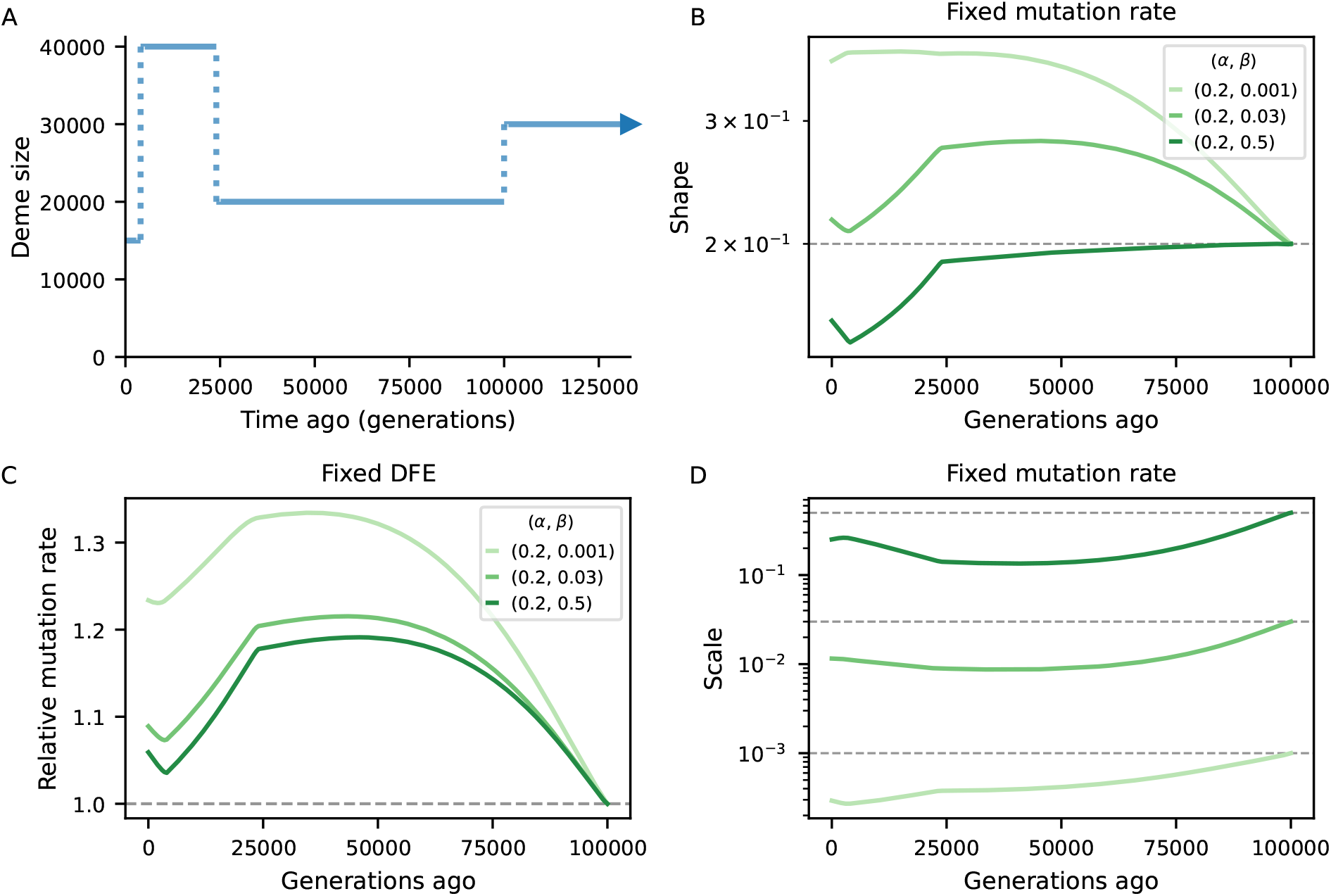
Biases in the inference of selection parameters due to the incorrect assumption of equilibrium. Population size trajectory (top left) is a rough piecewise constant representation of the Cousins et al. (2024) model. Other panels mirror Figure 4, except that shape and scale here combine for less deleterious DFEs. We either assume the mutation rate is known and fit the shape and scale of the DFE (panels B and D), or vice-versa (C).

Taken together, these results highlight the importance of jointly modeling demography and linked selection (Johri et al., 2020, 2023). The extent of the bias in *B*-maps and downstream inference will depend on the precise population history and DFE, but it can potentially be large. Previous studies may have misinterpreted patterns of diversity, potentially due to fitting probability weights to a discretized DFE which allowed too many degrees of freedom. Moreover, since both cBGS and the Buffalo and Kern (2024) implementation of SC16 show reduced accuracy around *N*_*e*_*s* = −1, our assessment of the temporal bias – arriving from `moments++` predictions – is conservative relative to what we may expect in previous studies.

## Conclusions

Accurately characterizing the effect of linked (negative) selection has been a goal of population genetics for decades (Charlesworth et al., 1993; Charlesworth, 2013; Cutter and Payseur, 2013; Charlesworth and Jensen, 2021). Pioneering studies focused on strongly selected sites (Charlesworth et al., 1993; Hudson and Kaplan, 1995b,a; Nordborg et al., 1996) while a subsequent framework managed to incorporate weak selection (Santiago and Caballero, 1995, 1998, 2016; Buffalo and Kern, 2024). These models have enjoyed success in predicting *π* across the genome, but their limitations in the interference selection regime and their assumption of equilibrium demography raise questions about their validity. Here we incorporated negative selection in the two-locus system (Hill and Robertson, 1968; Ragsdale and Gravel, 2019), which jointly models the evolutionary forces of drift, mutation, recombination and selection. The flexibility of such two-locus methods supports population-size history and a wide range of selection coefficients, which critically includes the region around *N*_*e*_*s* = −1, improving accuracy in several evolutionary scenarios while remaining computationally tractable. We also accommodated the Nordborg (1997) model to historical changes in population size, bridging the gap in *B*-value prediction, from weak to strong selection, for non-equilibrium populations. Exploring our new model, we found rich temporal dynamics of linked selection that depend on the DFE, genome structure, and precise demographic history.

Being able to readily compute expectations under arbitrary single-population scenario brings several benefits. For example, we showed that the temporal dynamics of *B*-values is highly dependent on the strength of selection and the prescribed demography (Figure S9). This clarifies the patterns highlighted by Figure S7 in Torres et al. (2020), where the trend may sometimes be hidden behind simulation noise. Instead, our approach provides expectations for diversity statistics in scenarios that would be computationally burdensome to extract from forward-in-time simulation (Figure S10). We note that Johri et al. (2021) derived expectations of *B*-values under two-epoch scenarios and assuming that *N*_*e*_*s, N*_*e*_*µ* and *N*_*e*_*r* remain constant (which holds as long as sampling takes place shortly after the population size change). They used these expectations to investigate the bias in demographic inference incurred by linked selection. Our more general model can foster the development of joint-inference approaches to address this important issue (Johri et al., 2020, 2023; Mackintosh et al., 2025).

Still there are caveats to our implementation. One technicality is that strong selection requires high orders of (1 − 2*p*)^*j*^ factors, which places the transition matrix close to singularity such that numerical instability may prevent the linear algebra algorithm from finding the steady-state of **v** (Figure 1). The critical point of instability is a function of *N* − (*t*), *s* and the order of 1 − 2*p*, which makes it difficult to propose a useful rule of thumb to circumvent it. In limited testing, we were always able to find an order of 1 − 2*p* that both displays high accuracy and prevents descent into numerical chaos. In practice, however, it is easier to simply use `moments++` in scenarios where |*s*| ≤ 0.005 and employ our extension of cBGS in the strong selection regime. Similarly, drastic demographic fluctuations can generate expectations of high-order moments of 1 − 2*p* that are numerically indistinguishable from zero in epochs of low *N*_*e*_. In such cases, scaling up the mutation rate restores numerical precision and is inconsequential to the prediction of *B*-values, which measure relative diversity.

A more conceptual challenge is in working out the interference correction under non-equilibrium demography, because population sizes fluctuations cause the dynamics of linked selection to change over time and thus the level of interference. Therefore, no single rescaling of local *N*_*e*_ can properly represent the historical linked selection exerted on a constrained site (analogously to neutral sites) and our approximate correction becomes improper. To circumvent this problem, in this article we focused on the temporal dynamics of *B*-maps in scenarios with relatively weak interference. We caution that assembling *B*-maps in species that experience severe size changes and widespread interference should require adjustments to take both into account.

Another current limitation of `moments++` is the restriction to single-population models. If non-equilibrium demography already creates complex temporal dynamics under panmixia, scenarios involving structure, migration and admixture may further exacerbate them (Hasan and Whitlock, 2024). Work is ongoing to assimilate gene-flow into the system and characterize these patterns. Finally, although `moments++` incorporates all two-locus statistics from the original basis **v**, here we only examined *π*_*R*_ (and *B*-values). Describing the behavior of LD statistics under linked selection is of interest, but we leave it for a future treatment.

Our main practical result is that inaccurate prediction of *B*-maps (either by neglecting a critical part of the DFE, assuming equilibrium demography, or both) biases downstream analyses (Figures 4, 5, S6). This helps explain why *B*-map based inferences of DFEs and deleterious mutation rates (McVicker et al., 2009; Murphy et al., 2022; Buffalo and Kern, 2024) produced estimates that are incompatible with other studies, although other factors could also be in play. These models were justified by their high predictive power, as measured by the variance in observed diversity explained by their *B*-maps, but aiming highly parameterized DFE models at the *B*-value landscape alone likely led to over-fitting. Although beyond the scope of this paper, work is underway to investigate models of human genetic diversity with moments++.

Besides DFE inference, *B*-values have also been used to investigate the impact of linked selection on the heritability of complex diseases, following the rationale that causal variants segregate more freely where the local effect of genetic drift is greater (Pardiñas et al., 2018; Li and Berg, 2025). In a similar vein, Simon and Coop (2024) showed that in recent European history, most of the temporal covariance in allele frequency change that would be attributable to linked selection can instead be explained by gene-flow, except in the lowest *B*-value bins. *B*-values are also routinely adopted in genomic masks, as a way to mitigate the background selection bias in demographic inference, e.g. Medina-Muñoz et al. (2023). We expect that our model will help to refine future analyses of this kind, especially when interest lies in close proximity to (weakly) constrained elements, where the cBGS theory employed so far struggles the most (Figure S6).

## Methods

### Model implementation

Due to the computational burden imposed by selection, we re-implemented the moments.LD framework (Jouganous et al., 2017; Ragsdale and Gravel, 2019) in C++. In doing so, we made two changes to the basic layout of the model.

First, we obtain the combined transition matrix *M* by multiplying together (rather than summing) the matrices that represent the individual operators (Selection, Recombination, Mutation, Drift, Figure 1C): *M* = *S* × *R* × *U* × *D*. This means that terms as small as 𝒪(*µrs/*2*N*) can be retained within *M*, rendering it slightly denser than the transition matrix in `moments.LD`. In practice, these inclusions have a negligible effect on the expectations such that predictions from `moments++` and `moments.LD` are indistinguishable under neutrality. Related, note that re-arranging the multiplication above amounts to changing the order of events in the life-cycle (drift represents reproduction), but this also has minimal impact on the predictions.

Second, `moments++` approaches demography as a collection of epochs with piecewise constant population sizes (Figure 1D), instead of the continuous-time treatment of `moments.LD`. This simplification brings about further computational efficiency and streamlines the design of evolutionary models to be explored in future studies, e.g. when selection coefficients, mutation rates and/or recombination rates change over time.

We benchmarked our model against moments.TwoLocus (Ragsdale, 2022) for different selection coefficients and orders of 1 − 2*p* factors (Figure S7). As mentioned in the concluding remarks, some parameter combinations may introduce numerical instability if the order of 1 − 2*p* factors is exceedingly high (e.g., when over-shooting it to guarantee accuracy in strong selection scenarios). In such cases, moments++ will throw an error and suggest adjusting this number.

### Extending cBGS to piecewise-constant histories

The expected reduction in diversity due to linked selection can be expressed as the expected reduction in the expected coalescence time (*T*_*MRCA*_) of a pair of samples experiencing linked selection relative to that under neutrality. In a constant-sized population, 𝔼 [*T*_*MRCA*_] = 2*N*_*e*_ under neutrality. Nordborg (1997) developed a structured coalescence model to obtain the expected *T*_*MRCA*_ given strong purifying selection at a linked locus, with mutation rate *u*, selection coefficient *s*, and genetic distance *r* between the selected and neutral loci.

We consider sampling two haplotypes, which could be found in the following states: both haplotypes are free of the deleterious allele (denoted (2, 0)), one carries the deleterious allele (denoted (1, 1)), or both carry the deleterious allele (denoted (0, 2)). The absorbing states are reached by coalescence from states (2, 0) and (0, 2), and are denoted (1, 0) and (0, 1). The transition matrix between states is given by Nordborg (1997) (Equation 2) as

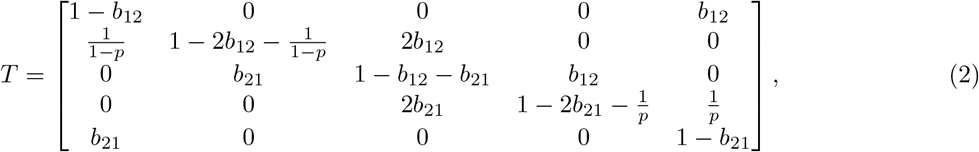

where *p* = *u/s* is given by mutation-selection balance, and the coefficients *b*_12_ and *b*_21_ are

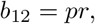

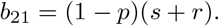

Nordborg (1997) showed that the total coalescence rate in this model is given by

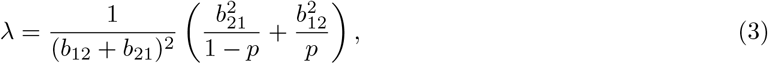

which is greater than one, so that

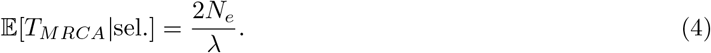

To account for piecewise-constant population size changes, we consider the probability that coalescence occurs within a given epoch and compute the *T*_*MRCA*_ conditioned on coalescence occurring within that epoch. In this case,

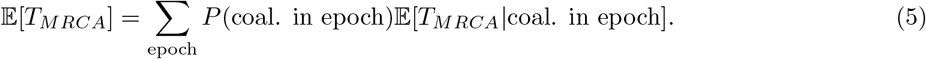

For the neutral coalescent,

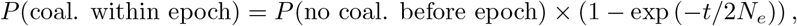

where the length of the epoch *t* is measured in generations. The probability that no coalescence occurs before a given epoch can be iteratively obtained over the preceding epochs, looking backward in time. In the model with selection, this can be found using the sub-intensity matrix of the transition matrix (Equation 2),

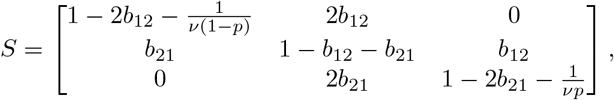

in which 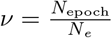. Here, *N*_*e*_ is taken to be some reference population size, often arbitrarily chosen as the size in the most ancient epoch. Using the expected relative frequencies of occupying each transient state under mutation-selection balance, we then have

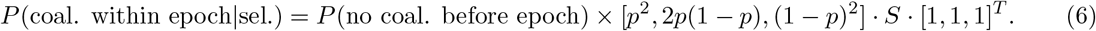

The expected coalescence time, conditioned on coalescing within an epoch, can be found using the coalescence rate *λ* (Equation 3). In an epoch spanning time [0, *t*),

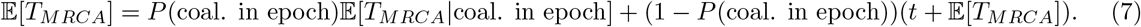

Noting that 𝔼[*T*_*MRCA*_] = 2*N*_*e*_*/λ* and rearranging, we find

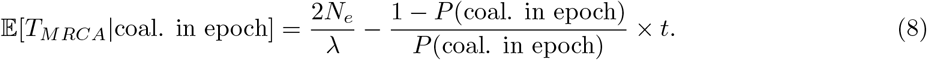

Then for any epoch spanning time [*t*_0_, *t*_*f*_), we can use this same formula, with *t* = *t*_*f*_ − *f*_0_, and simply add *t*_0_to account for the translation of time to the start of the given epoch.

### Multi-locus prediction

At its heart, `moments++` is a pure two-locus model in which the focal neutral site is linked to a single constrained site at arbitrary genetic distance (Figure 1A). Our approach to predicting *B*-values in the presence of several constrained elements involves pre-computing a lookup table that stores, for a prescribed demographic model, pure two-locus predictions on a parameter grid. In this article, each of the lookup tables built with `moments++` embodied 35 selection coefficients (from 0 to −0.001) and 72 recombination rates (from 0 to 0.5), roughly linearly spaced in logarithmic scale. We then used our extension of the Nordborg (1997) model to incorporate more deleterious selection coefficients (down to −1), and performed cubic splines interpolation to have a smooth function of *B* against *r* that is easily applicable along the chromosome. When elements were constituted by contiguous sites, we computed the *B*-value reduction by multiplying together the effect of each individual site (i.e., we assume no within-element interference). In doing so, we considered the genetic distance between our focal site and the midpoint of the element. Both approximations (no interference and midpoint recombination distance) are fairly accurate for elements with ~ 1000 sites or less. When dealing with chromosome layouts that mimic the position and lengths of exons in humans, we divided elements with > 1000 sites into smaller adjacent elements. In models where selection coefficients follow a continuous DFE, we used numerical integration to obtain probability weights at each of the selection coefficients stored in the lookup table. In such cases, individual exons were assumed to follow the same DFE and therefore shared probability weights.

Lookup tables contain a single mutation rate (in this article, *µ* = 10^−8^ per site per generation). When relevant (e.g., performing interference correction), we applied a linear adjustment of the *B*-value with respect to element-specific mutation rates. Incorporating a heterogeneous mutation landscape becomes straightforward in this setting, although we did not explore such scenarios here.

### Interference correction

Selective interference can be thought of as linked selection among constrained sites. Borrowing from cBGS (Charlesworth et al., 1993; Charlesworth, 2009), we developed a correction scheme where constrained elements impose reductions in each other’s effective population size (Good et al., 2014). The reduction in *N*_*e*_ of a given element is then equivalent to its *B*-value. To mimic this effect, at each round of prediction we scale *µ, r* and *s* of every element by its corresponding *B*-value. The justification is that the dynamics are determined by the scaled rates *N*_*e*_*µ, N*_*e*_*r* and *N*_*e*_*s*. In turn, this scaling changes the linked selection effect that the elements exert on each other (and on neutral sites), thus we re-compute *B*-values all around. By iterating this procedure, we can obtain a *B*-map that approximately accounts for interference among constrained sites. In practice, this converges in < 10 iterations (often fewer), even for dense regions with strong linked selection effects (Figure 2, S1).

### *B*-map predictions

The chromosome layout used to assess the bias in DFE inference under non-equilibrium (Figure 4, 5) used *µ* = 10^−8^, *r* = 10^−8^ as well as an arbitrary 10 Mb chromosome where 20 functional elements, each with length 1 kb, are evenly distributed. Within each element, the nonsynonymous (constrained) to synonymous (neutral) ratio was set to 2.31: 1 (Kim et al., 2017). The DFE parameters varied and are depicted in Figures 4, 5.

When assessing the bias incurred by incorrectly assuming equilibrium, the *N*_*e*_ used to predict equilibrium *B*-maps was based on the expected diversity coming from a purely neutral model (*N*_*e*_(*t*) = *π*_0_(*t*)*/*4*µ*). Here, *π*_0_(*t*) was extracted from moments++ predictions of a neutral model that follows the same demography as the linked selection model to which it is being contrasted.

### Forward simulations

Multi-locus forward-in-time simulations (Figure 2, 3) were conducted using fwdpy11 (Thornton, 2014, 2019). We computed pairwise diversity at each site (and by extension, *B*-values) from the tree sequences using the expected number of mutations given the realized branch lengths (removing mutational noise) and averaged over 10,000 replicates to reduce genealogical noise. Thus, our goal was not to ask how much of the variance in diversity is captured by the predictions, but to quantify the bias in moments++ using a low-variance estimate of *π*_*R*_.

In benchmarking simulations (Figure 2), per-base recombination and mutation rates were uniform, with *r* = 10^−8^ and the deleterious rate *µ* taking values between 10^−9^ and 10^−8^. Population sizes were held constant at 10^4^ diploid individuals. In simulations following human genomic annotations, the recombination rates were determined by the OMNI recombination map inferred from YRI individuals in the Thousand Genomes Project (Consortium et al., 2012). Mutation rates were held constant at *µ* = 1.5 × 10^−8^, with the nonsynonymous (selected) mutation rate within annotated exons equal to 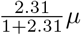 (as the ratio of new mutations within exons being nonsynonymous:synonymous has been estimated to be 2.31: 1 in humans (Huber et al., 2017)). Coding regions were specified by annotated exons in the human genome, using human genome build 37. Selection coefficients were drawn from a gamma-distributed distribution of fitness effects, with shape parameter 0.215 and scale parameter 0.028105 (Kim et al., 2017).

We also developed `twoLocusSim`, a forward-in-time simulator of independent two-locus systems written in `C++`. Briefly, it employs an infinite-sites mutation model to spawn one- and two-locus diversity with a pseudo-random number generator. Recombination and selection operate deterministically on haplotype frequencies, and drift is implemented through multinomial sampling of haplotypes every generation. The software can output statistics in time-series fashion, and we used `twoLocusSim` to understand the mechanism of selection in the Hill-Robertson system (Appendix B).

### Data and software availability

Source code for `moments++` and `twoLocusSim` is available at https://github.com/gvbarroso/momentspp (Barroso and Ragsdale, 2025). The version of the `moments++` source code used to generate the results in the present article has been frozen and stored in https://doi.org/10.5281/zenodo.17136261. High-level tools for predicting *B*-maps (using a lookup table that stores moments++ predictions) are available as a python package called `bgshr`, available at https://github.com/apragsdale/bgshr (Barroso et al., 2025). The version of `bgshr` used to generate the results in the present article has been frozen and stored in https://doi.org/10.5281/zenodo.15802044. Jupyter notebooks used to generate figures, as well as the associated metadata, can be found at https://github.com/gvbarroso/momentspp/tree/main/doc/examples/figures_paper1. Scripts used to perform simulations are provided in https://github.com/gvbarroso/momentspp/tree/main/doc/examples/figures_paper1/fwdpy11_simulations. The Jupyter notebooks and `fwdpy11` simulation scripts are packaged together with the `moments++` source code (https://doi.org/10.5281/zenodo.15824891).

## Acknowledgments

We thank members of the Ragsdale lab for helpful discussions on this work, Nick Collier for careful comments on the manuscript, and Lloyd Kirk for implementing the Docker container build. This work was supported by NIH Award R35-GM154962 to APR.

## Appendix

### A The two-locus system with selection

By relating two-locus statistics to the evolutionary processes that affect them, Hill and Robertson (1968); Ohta and Kimura (1969) developed a system of equations for studying LD under neutrality. They found that solving for the evolution of 𝔼 [*D*^2^] under drift and recombination (where *D* is the standard covariance measure of LD) requires the inclusion of 𝔼 [*D*(1 − 2*p*)(1 − 2*q*)] and 𝔼 [*p*(1 − *p*)*q*(1 − *q*)] (*p* and *q* being the derived allele frequencies at the left and right loci, respectively). With mutation, the expected pairwise diversity at each locus is needed, that is, 𝔼 [2*p*(1 − *p*)] and 𝔼 [2*q*(1 − *q*)]. Together, this set of summary statistics, which we denote **v**, is closed under neutrality. That is, a full description of their dynamics depends only on other statistics already present in **v**, leading to a finite number of recursions (Figure 1).

We may think of these statistics (or moments) as representing sampling configurations of four two-locus haplotypes (Ragsdale and Gravel, 2019; Ragsdale, 2022). From this angle, the individual effects of evolutionary processes become apparent. Mutation introduces one-locus diversity in the system by creating a derived allele in a previously monomorphic sample; similarly, it introduces two-locus diversity by creating a derived allele in the monomorphic locus of a sample that already segregates at the other. Recombination shuffles alleles between the two haplotypes, whereas drift reduces both one- and two-locus diversity when a haplotype copies over another (backward-in-time, a coalescence event). This perspective aids modeling other processes such as migration and admixture (Ragsdale and Gravel, 2019; Ragsdale et al., 2023). We now want to incorporate selection into the Hill-Robertson system.

The effect of negative selection on the two-locus sampling configurations is to remove a haplotype carrying the deleterious variant with rate proportional to its fitness effect (in diploids, this is equivalent to semi-dominant selection acting at the gametic stage). Under a soft selection model (i.e., selection does not alter population sizes), the eliminated haplotype must be replaced by a haplotype randomly chosen from the rest of the population. We derive the selection dynamics in expectation, reaching a deterministic model where selection steadily removes a fraction of haplotypes carrying deleterious variants from the population at each generation. Denoting the pair of ancestral and derived alleles at the left locus (constrained) by *a, A* and similarly *b, B* for the right locus (neutral), the expected haplotype frequencies after one generation of selection are (to first order in *s*):

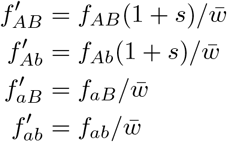

with 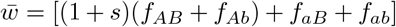 the mean fitness of the population (Ragsdale and Gravel, 2019, see section S1.1.4 of the Supplement in).

Converting from two-locus haplotypes to the space of statistics where the Hill-Robertson basis resides, negative selection directly reduces diversity at the left (constrained) locus, as

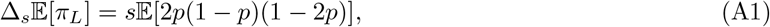

where *π*_*L*_ = 2*p*(1 − *p*). Since *s* is negative and 𝔼[2*p*(1 − *p*)(1 − 2*p*)] is positive, 𝔼[*π*_*L*_] decreases due to direct selection. Moreover, the dependence of *π*_*L*_ on an additional statistic (*π*_*L*_(1 − 2*p*)) immediately hints at the non-closure of the system under selection (Jouganous et al., 2017). On the other hand, selection *indirectly* reduces diversity at the neutral locus (equation 1, Figure 1).

The mechanism of haplotype elimination and replacement creates an intricate pattern of dependence among the terms in **v**. When one of the two loci is constrained while the other remains neutral, the ancestral-derived symmetry in the constrained locus is broken, as well as the left-right symmetry between loci, and the system no longer closes. This means that in the presence of selection, expectations of two-locus statistics in the next generation depend on expectations of statistics that are absent from the original basis. These, in turn, need additional moments themselves, so the system grows indefinitely. Intuitively, such growth could be anticipated from the way selection operates: to replace the eliminated haplotype, we need to know the expected frequency of haplotypes in a larger sample (Jouganous et al., 2017). Thus the expectation of, e.g., *D* after one generation of selection requires contributions of statistics of the form *D*(1 − 2*p*)^*j*^ and so on, recursively, where the order *j* is unbounded (Figure S2). A graphical representation of the selection operator acting on 𝔼 [*D*] and extending to the first three layers of extra moments is given in Figure S2.

### B Proximate and ultimate explanations for the decay of *π*_*R*_ in the Hill-Robertson system

The goal of this section is to understand how negative selection operates in (*p, q, D*)-space (Ragsdale and Gravel, 2019). To this end, it is instructive to discuss why 𝔼[*D*(1 − 2*q*)] should be positive in the presence of selection. With hindsight, this must be true otherwise *π*_*R*_ would not decay through linkage with a selected locus (Equation 1). Without this retroactive argument, however, the reason is not obvious. *D*(1 − 2*q*) is a LD statistic that receives greater contribution from variants at the tails of the neutral locus’ frequency spectrum, with change between consecutive generations given by

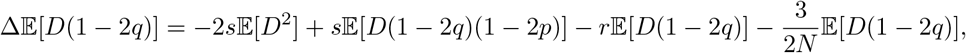

where the first two terms on the right-hand side embody the selection operator (*s* < 0). Its neutral expectation is zero, simply decaying by recombination and drift, but under negative selection 𝔼[*D*(1 − 2*q*)] > 0 indicates that the buildup through *D*^2^ is higher than the decay through 𝔼[*D*(1 − 2*p*)(1 − 2*q*)]. Why should this be?

Taking a step back, we note that 𝔼[*D*] equals zero under both neutrality and selection. To understand why, first note that *D* = 0 until two-locus diversity emerges. This second mutation event will either create “coupling” (*AB*, ⇒ *D* > 0) or “repulsion” (*aB* or *Ab*, ⇒ *D* < 0) haplotypes. From the infinite sites model and the assumption of weak mutation (*N*_*e*_*µ* ≪ 1), it follows that 𝔼[*p*] < 1*/*2 and 𝔼[*q*] < 1*/*2. That is, single-locus diversity implies a derived allele is expected to segregate at low frequency, regardless of which locus mutates first. Therefore, the second mutation event in the two-locus system is more likely to produce “repulsion” haplotypes. Naively, this would lead us to believe that mutation tends to push *D* < 0, but this higher probability is balanced by the increased value of *D* > 0 whenever mutation does generate “coupling” haplotypes. Hence mutation does not affect *D* in expectation (Ragsdale and Gravel, 2019). Meanwhile, recombination and drift lead to its decay:

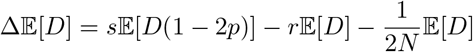

Here we see that selection on *D* collects only from *D*(1 − 2*p*), which itself evolves according to

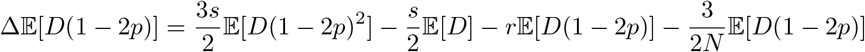

and the recursions for higher-order 𝔼[*D*(1 − 2*p*)^*j*^] moments follow the same pattern of exclusive contributions from the adjacent 𝔼[*D*(1 − 2*p*)^*j* − 1^] and 𝔼[*D*(1 − 2*p*)^*j*+1^] through selection:

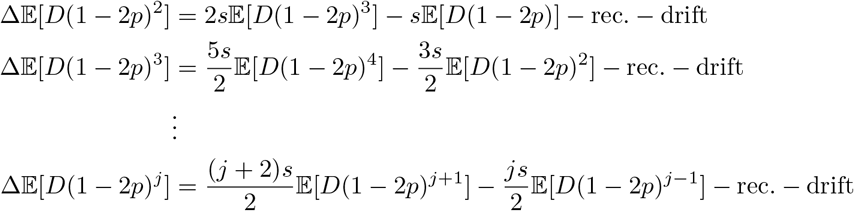

`moments++` reveals that all 𝔼 [*D*(1 − 2*p*)^*j*^] moments are zero regardless of selective constraint. When *s* < 0, each collects from their adjacent neighbors with opposite signs and in the end these contributions cancel out. This means that 𝔼 [*D*] is flat over the distribution of derived allele frequencies at the constrained locus (Figure A1A). Indeed, the change in *p* after one generation of selection,

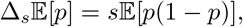

is independent of the neutral locus. Therefore conditioning on *p* (via 1 − 2*p*) should not affect the covariance between allele frequencies. Conversely, 𝔼[*D*(1 − 2*q*)] > 0 suggests that 𝔼[*D*] is *not* flat over the distribution of derived allele frequencies at the neutral locus (Figure A1B). This happens because the expected change in *q* after one generation of selection,

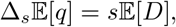

is entirely attributed to linkage with the constrained locus. (Hence conditioning on *q* (via 1 − 2*q*) does affect the expectation of the covariance term *D*.) In fact, since 𝔼 [*D*] is zero, the two-locus system predicts that selection does not affect *q* in expectation. This happens because the decrease in frequency when the neutral mutation falls on the derived (deleterious) background is compensated by its increase when it falls on the ancestral background, which is relatively more fit. (A similar logic applies when the neutral locus segregates first.) In other words, linked selection should not change the mean of the neutral allele frequency spectrum, instead distorting its outline by increasing mass at both tails (Ewing and Jensen, 2016; Cvijović et al., 2018) – in line with the classic result that linked selection does not alter the rate of fixation of neutral alleles (Birky Jr and Walsh, 1988). This formulation is diametrically opposed from cBGS models which postulate that the reduction in *π*_*R*_ comes from a decrease in *N*_*e*_, changing the mean of the neutral allele frequency spectrum without touching its outline.

Having built some intuition, we come back to the original question: how does negative selection cause 𝔼[*D*(1 − 2*q*)] > 0? Two alternative mechanisms could be in play. First, this distortion could be manifested immediately upon the mutation event that generates two-locus diversity. In this case, selection would influence 𝔼[*D*(1 − 2*q*)] through its “stationary” effect on the site frequency spectrum. Second, even if 𝔼[*D*(1 − 2*q*)] = 0 immediately after the second mutation, selection could distort its expectation over the course of time. This could happen if elimination of haplotypes carrying the deleterious allele does not preserve the symmetry of *D* (the canceling out of negative and positive instances) over the distribution *f* (*q*).

We tested whether 𝔼[*D*(1 − 2*q*)] > 0 upon the the second mutation event (first hypothesis). Conditioning on the right locus segregating first, then prior to the second mutation we have *f*_*aB*_ = *q* and *f*_*ab*_ = 1 − *q*.

With probability *q*, the second mutation creates the *AB* haplotype at frequency 1*/*2*N*, in which case *D* = *f*_*ab*_*f*_*AB*_ − *f*_*aB*_*f*_*Ab*_ = (1 − *q*)*/*2*N*. Otherwise, (probability 1 − *q*) it creates the *Ab* haplotype and *D* = −*q/*2*N*. Weighing *D*(1 − 2*q*) in both cases by their respective probabilities and integrating over the neutral frequency spectrum *f* (*q*) ∝ 1*/q*, we have

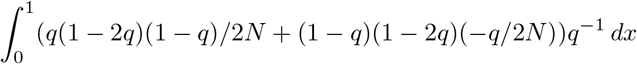

which yields zero, therefore, immediately after the second mutation event and if the right locus segregate first, 𝔼 [*D*(1 − 2*q*)] = 0. Similarly, if the left locus segregates first we integrate over the deleterious site frequency spectrum *f* (*p*) (Ewens, 1964) to obtain

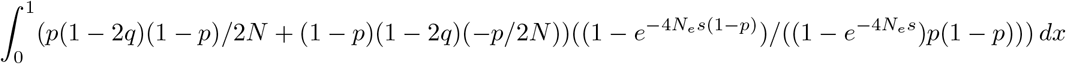

which likewise yields zero, thus we reject the first hypothesis. We conclude that selection builds up 𝔼 [*D*(1 − 2*q*)] over the course of time.

To understand why, consider again the situation where the right locus segregates first. We have seen that, immediately after the left mutation, 𝔼 [*D*] = 0 regardless of *q*. However, if *q* < 1*/*2, then selection will mostly act against *Ab* haplotypes, such that over time *D* will tend to be positive on the first half of the neutral frequency spectrum. Conversely, if *q* > 1*/*2, then selection will mostly act against *AB* haplotypes, such that over time *D* will tend to be negative on the second half of the neutral frequency spectrum. Both situations lead to 𝔼 [*D*(1 − 2*q*)] > 0 and we conclude that selection distorts the distribution of *D* over *f* (*q*) by virtue of its effect after two-locus diversity is established.

We confirm these trends in forward-in-time simulations with twoLocusSim (Figure A1). These simulations further allow us to pinpoint thar *D* increases at first, then decreases, whereas 𝔼 [*D*(1 − 2*q*)] tends to remain above zero over time (Figure A2).

### C Temporal dynamics in the pure two-locus model

We now discuss the behavior of *π*_*R*_ when the focal neutral site is linked to a single constrained site. For simplicity, we describe how *B*-values evolve in two-epoch demographic models (10-fold bottleneck or expansion). Patterns from this pure two-locus model will help us interpret multi-locus scenarios (Figure 2, 3).

A comprehensive description of the temporal dynamics at the neutral locus requires understanding the behavior of the linked constrained locus. Therefore, we start by analyzing the trajectory of *π*_*L*_ as a function of its selection coefficient and demographic model. This corresponds to a single-site model whose behavior was described by Otto and Whitlock (1997) and has led to DFE inference methods based on the frequency spectrum of (independent) constrained sites (Boyko et al., 2008; Keightley and Eyre-Walker, 2007; Kim et al., 2017). Although not novel, the prediction of *π*_*L*_ is part of the `moments++` framework and we revisit it here to aid the interpretation of *B*-value dynamics.

We track the expected value of *π*_*L*_ for 120,000 generations following a population size change. As expected, the patterns are markedly different between bottleneck and expansion scenarios (Figure S8). Focusing on the expansion first, we see that if selection in the ancestral population is strong (2*N*_*e*_*s* = −20, Figure S8A), then *π*_*L*_ increases ever so slightly at first but pulls back to the previous equilibrium point. This happens because the steady-state in the ancestral population is already dominated by mutation-selection balance (*π*_*L*_ ≈ *u/s*) such that a 10-fold increase in *N*_*e*_ can only move the population deeper into the same regime. On the other hand, if the ancestral population experiences weak-to-moderate selection (Figure S8B, 2*N*_*e*_*s* = −2), a 10-fold expansion brings it close to mutation-selection-balance. After an initial uptick in *π*_*L*_ caused by the reduced coalescence rate, selection starts purging the excess of deleterious variants. Steady-state lies intermediate between ancestral *π*_*L*_ and such peak. Finally, if the ancestral population finds itself in the nearly-neutral regime (Figure S8F, 2*N*_*e*_*s* = −0.2), where the equilibrium *π*_*L*_ is relatively close to the neutral expectation (*π*_0_ = 4*N*_*e*_*µ*), a 10-fold expansion will introduce weak-to-moderate selection. *π*_*L*_ will rise smoothly, driven by the 10-fold increase in the scaled mutation rate, but eventually plateau at a point distinguishably different from *π*_0_, as selection becomes a relevant force.

The temporal dynamics of *π*_*L*_ are very different in the bottleneck case. If selection in the ancestral population is not strong (2*N*_*e*_*s* = −2 or 2*N*_*e*_*s* = −0.2, Figure S8B, C), *π*_*L*_ decays monotonically and quickly reaches the new equilibrium. However, if selection is strong enough (2*N*_*e*_*s* = −20, Figure S8A), a quick drop in *π*_*L*_ is followed by partial recovery towards an intermediate equilibrium point. This happens because the population suddenly leaves mutation-selection balance to enter the mutation-drift-selection regime. Deleterious mutations are now met with selection that is effectively weaker than in the ancestral population, thus *π*_*L*_ upticks after the initial drop. Note that in both demographic models the absolute change in *π*_*L*_ under strong selection (y-axes in Figure S8A) is quite low. In fact, in the expansion case the changes are negligible (as predicted by theory), however, in the bottleneck case the modest non-monotonic behavior of *π*_*L*_ promotes interesting dynamics of diversity at the linked neutral site, which we describe next.

**Figure A1:**
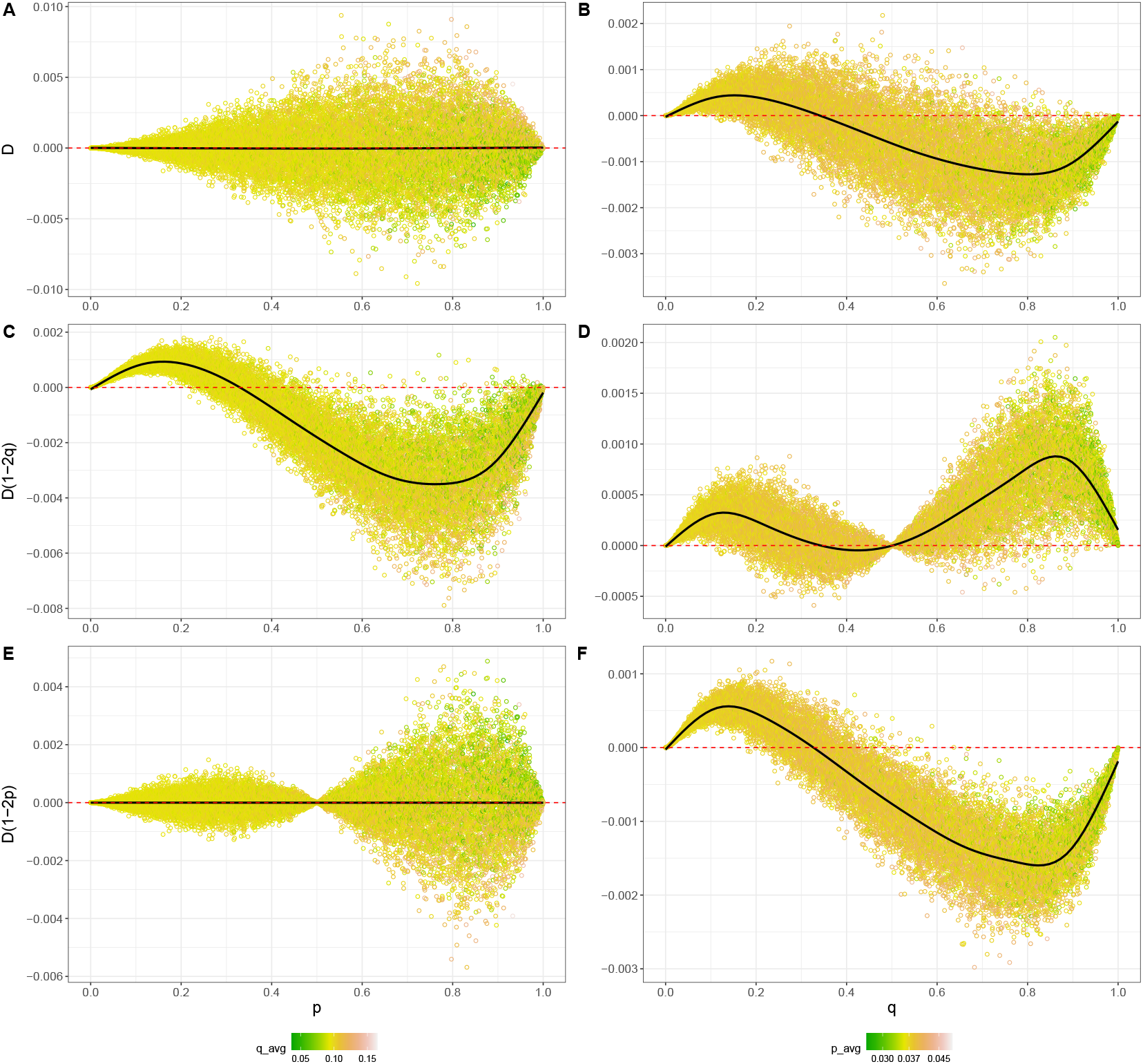
*D* statistics simulated with twoLocusSim. In each of 12,000 replicates, independent two-locus systems evolved for 10^7^ generations (sequence length *L* = 10) with *µ* = 10^−6^, *r* = 10^−4^ and *s* = −10^−4^. Each point represents the average over all generations (after a burn-in period of 400,000 generations), conditioning on a particular frequency of *p* (A, C, E) or *q* (B, D, F). Black line shows the loess smoothing curve. Note that *D* is flat (in expectation) over *f* (*p*) (A), but not over *f* (*q*) (B).

Population expansions lead to lower equilibrium *B*-values relative to the ancestral population (Figure S9, S10). When selection in the ancestral population is already strong (2*N*_*A*_*s* = −20), the *B*-value increases for many generations before starting to decline. This initial uptick means that the rate of change of *π*_*R*_ is, at first, faster than that of *π*_0_. Eventually, the *B*-value returns to the previous steady-state, as expected from cBGS predictions. On the other extreme, if deleterious variants behave nearly-neutrally in the ancestral population (2*N*_*e*_*s* = −0.2), a 10-fold expansion will send them to the weak-to-moderate regime. Driven by the increase in *N*_*e*_, overall levels of diversity increase, but the higher *π*_*L*_ renders the increase in *π*_*R*_ smaller than that of *π*_0_ and the *B*-value goes straight down. Finally, when 2*N*_*e*_*s* = −2 a 10-fold expansion imposes efficient selection. For the first ~*N*_*e*_ generations, the *B*-value curve resembles the mirror image of *π*_*L*_, decreasing at first and then creeping back up. Due to the increased deleterious diversity, the *B*-value then slowly drops below its ancestral steady-state. Overall, these dynamics are quantitatively dependent on the recombination rate, where strongly deleterious mutations reduce *B*-values over long genetic distances while weakly deleterious mutations have a much more localized effect.

**Figure A2:**
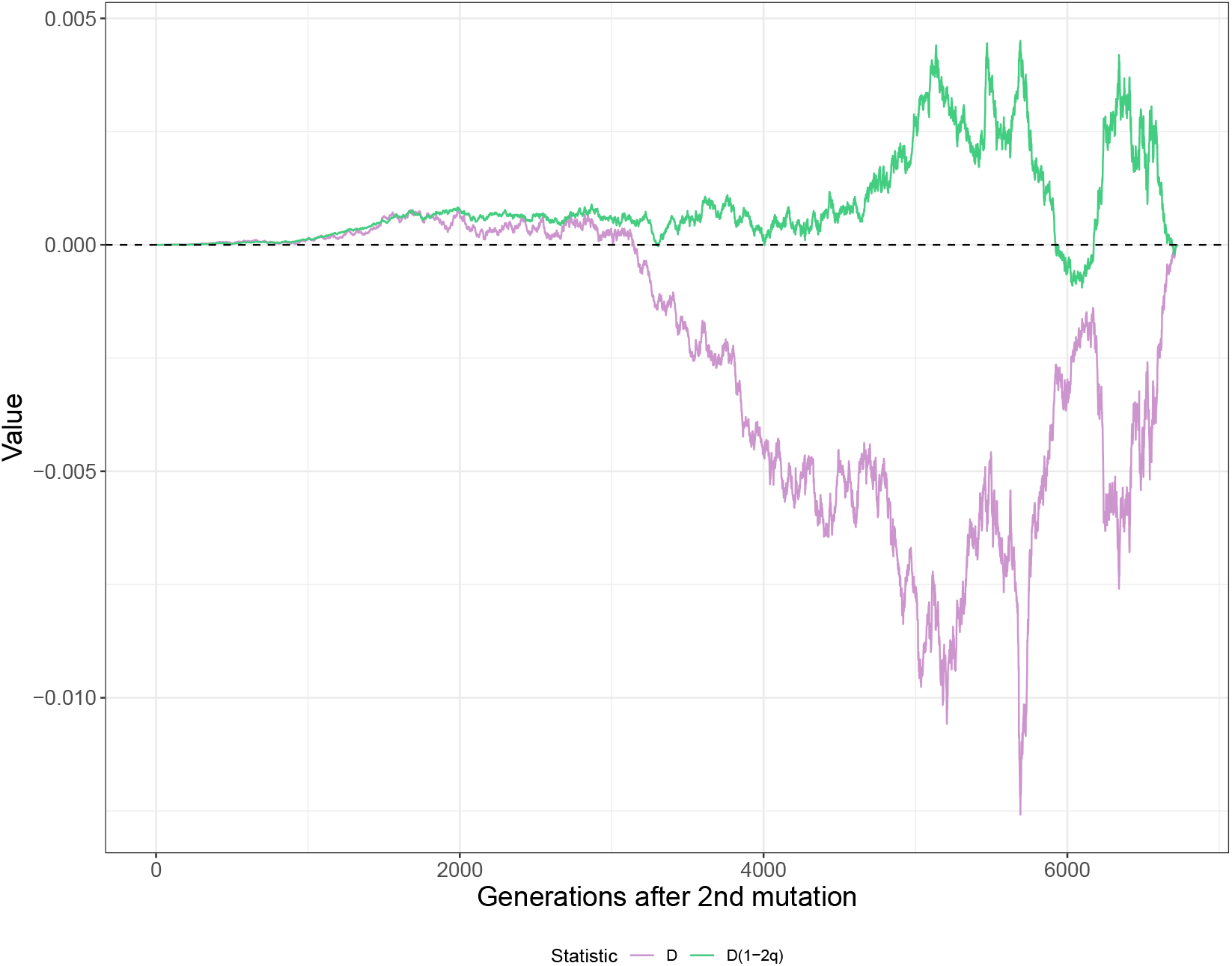
The temporal dynamics of *D* statistics after two-locus diversity is created. In each of 12,000 replicates, independent two-locus systems evolved for 10^9^ generations (sequence length *L* = 1) with *N*_*e*_ = 10, 000, *µ* = 10^−8^, *r* = 10^−8^ and *s* = −10^−3^. Shown are the average statistics over all replicates, conditioning on two-locus diversity reaching a particular generation (i.e., no fixation or loss). Therefore, since the number of simulations still segregating decreases over time, noise in these estimates increases and becomes substantial after ~ 5, 000 generations.

Population bottlenecks lead to higher equilibrium *B*-values relative to the ancestral population (Figure S9). Since these new equilibria do not depend on the slow supply of new mutations, they are attained much faster than in the expansion scenarios. When the ancestral population is under weak and moderate selection the increase is monotonic whereas the strong selection scenario reveals an initial drop in B, followed by recovery and beyond. In the latter case, the drop in *π*_*R*_ is initially more pronounced than in *π*_0_ because the approach to the new equilibrium happens faster in the presence of selection (see subsection “Modeling strong selection under non-equilibrium demography” and Torres et al. (2020)). Then, as *π*_*L*_ recovers but settles at an intermediate point (Figure S8), linked selection is relaxed relative to the ancestral population, and the *B*-value finds an elevated steady-state. In all selection regimes, the relationship between the equilibria points for *B* and *π*_*L*_ is intuitive: the first rises whenever the second drops. Under a 10-fold bottleneck, it is the stronger selection regime that causes non-monotonic dynamics. As in the expansion scenarios, recombination primarily modulates the magnitude of linked selection while preserving the shape of temporal dynamics.

The results from two-epoch models support that equilibrium *B*-values correlate negatively with *N*_*e*_ (Nordborg et al., 1996; Torres et al., 2020). (This explains why interference – approximated as a reduction in the *N*_*e*_ of constrained elements – always weakens linked selection effects). Indeed, this trend holds as long as the population is not under mutation-selection balance, in which case *π*_*L*_ is dictated by the ratio *µ/s* and independent of *N*_*e*_, as predicted by theory. The same logic applies under non-equilibrium demography, but the distinct evolutionary pace of *π*_0_, *π*_*R*_ and *π*_*L*_ shapes the temporal trajectory of *B*-values. Their transient behavior can be non-monotonic and last for several hundred generations, longer than most populations remain static in nature.

## Supplemental Figures

**Figure S1:**
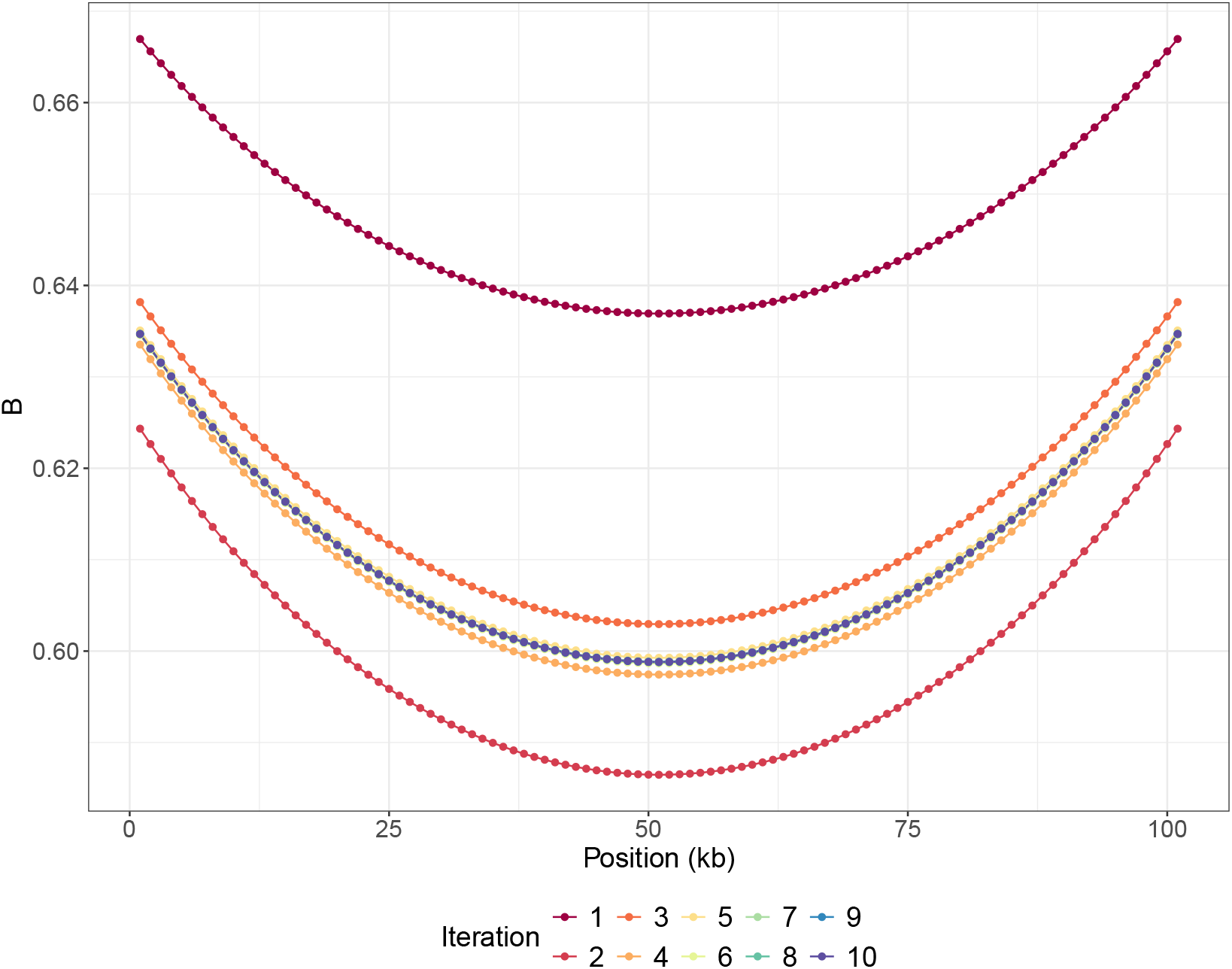
Predicted *B*-maps along a 100 *kb* segment, after each round of interference correction (color). Note how the iterative re-scaling of *µ, r*, and *s* leads to *B*-maps bouncing around the final values (convergence happens visually after the 6th iteration). Here the 100 kb segment layout follows that of Figure 2

**Figure S2:**
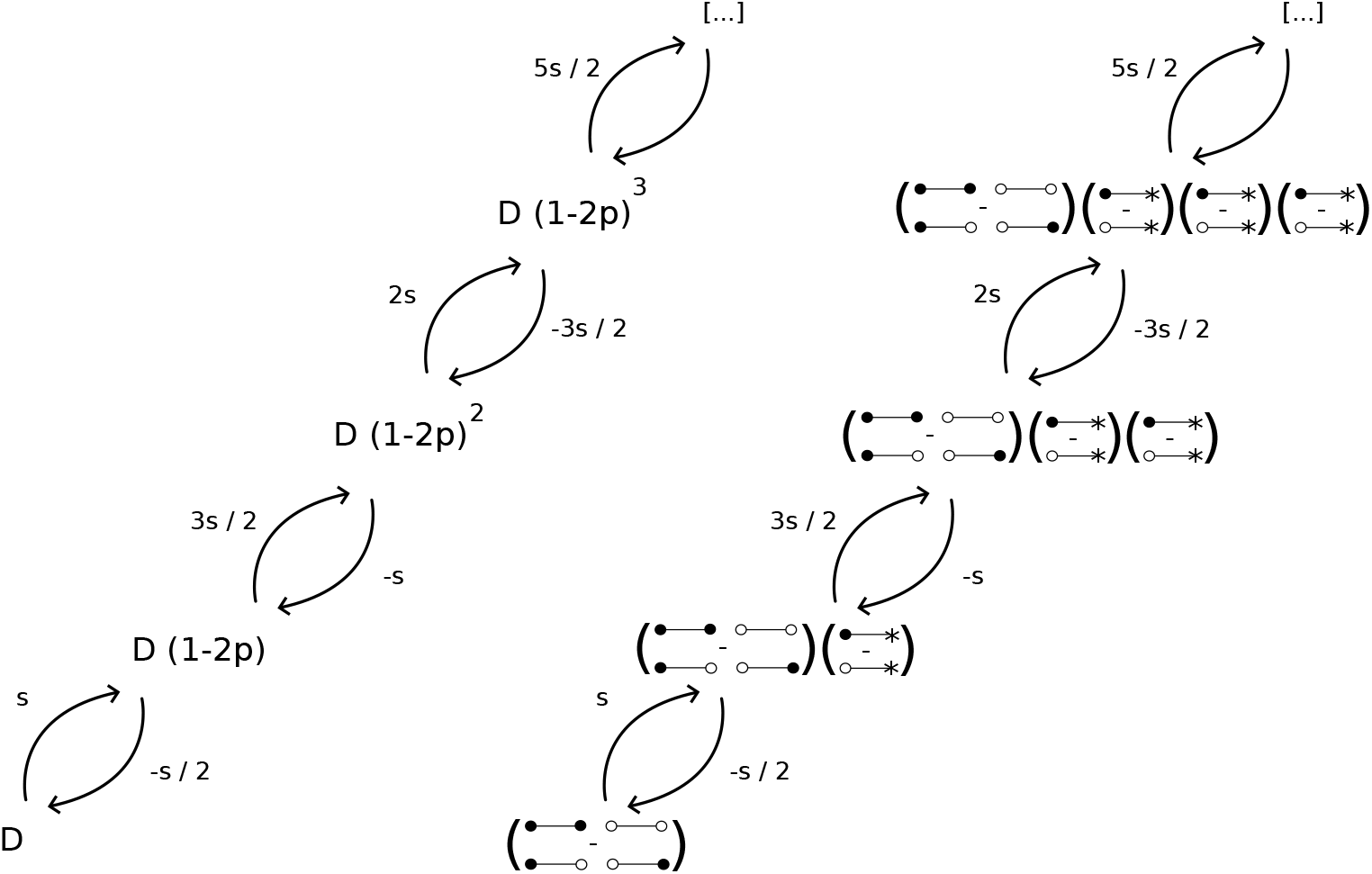
Representation of the dependencies among *D*(1 − 2*p*)^*j*^ statistics imposed by the selection operator in (*p, q, D*)-space (left) and the equivalent haplotype space (right). Arrows denote “collects from”, with the respective entry in the selection matrix shown above (*s* < 0). Haplotype subtractions proceed from top to bottom of the configuration within parentheses. Asterisks denote invariance to the allelic state at the right locus (1 − 2*p*).

**Figure S3:**
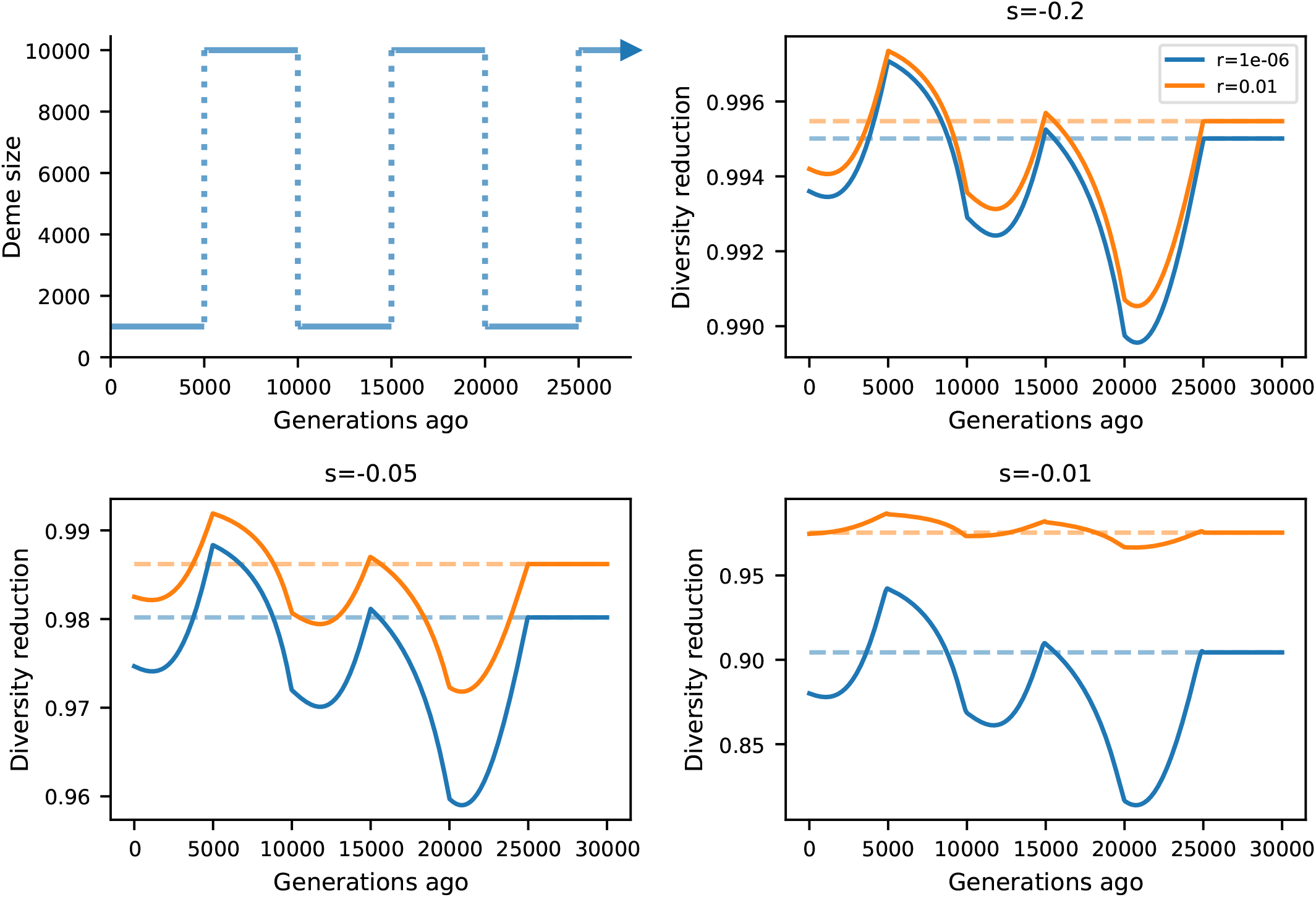
Temporal dynamics of BGS due to strong selection, as predicted using our extension to Nordborg (1997). Top left panel shows the demographic history considered. Each other panel shows, for a given selection coefficient, the predicted *B*-value trajectory for either tightly linked (*r* = 10^−6^, blue lines) and loosely linked loci (*r* = 10^−2^, orange lines). Dashed lines indicate the respective steady state solutions. Here the mutation rate is set to *u* = 10^−3^.

**Figure S4:**
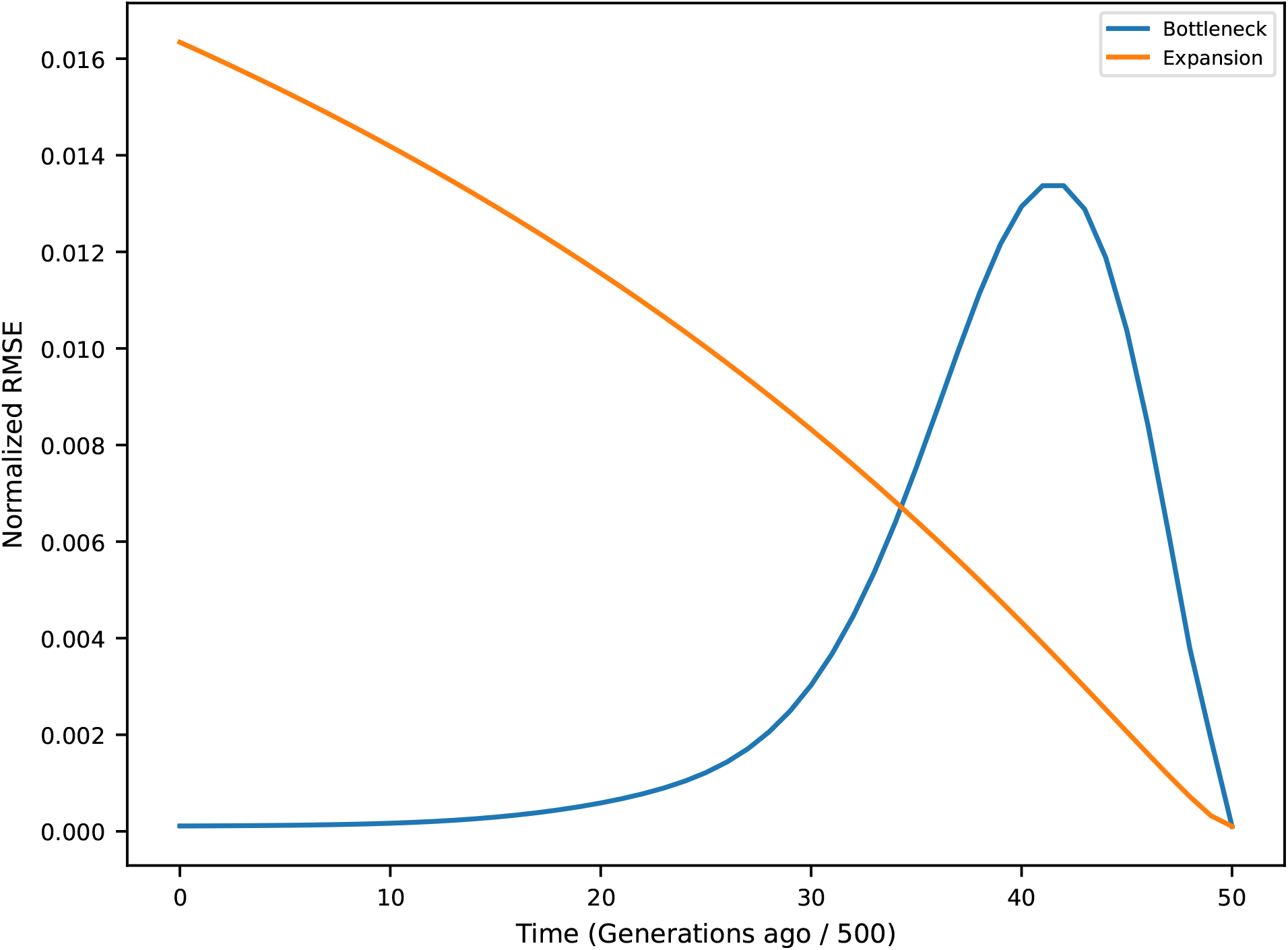
Temporal evolution of the the Normalized Rooted Mean Squared Error between the demography-aware *B*-map and steady-state *B*-maps, predicted every 500 generations after the size change. Blue and orange lines show results from a 10-fold bottleneck and expansion, respectively. Steady-state *B*-maps are obtained using the equilibrium *N*_*e*_ value calculated from neutral genetic diversity in the absence of linked selection (equivalent to the harmonic mean of population sizes in the relevant time-frame). Here the chromosome layout follows that of Figure 3.

**Figure S5:**
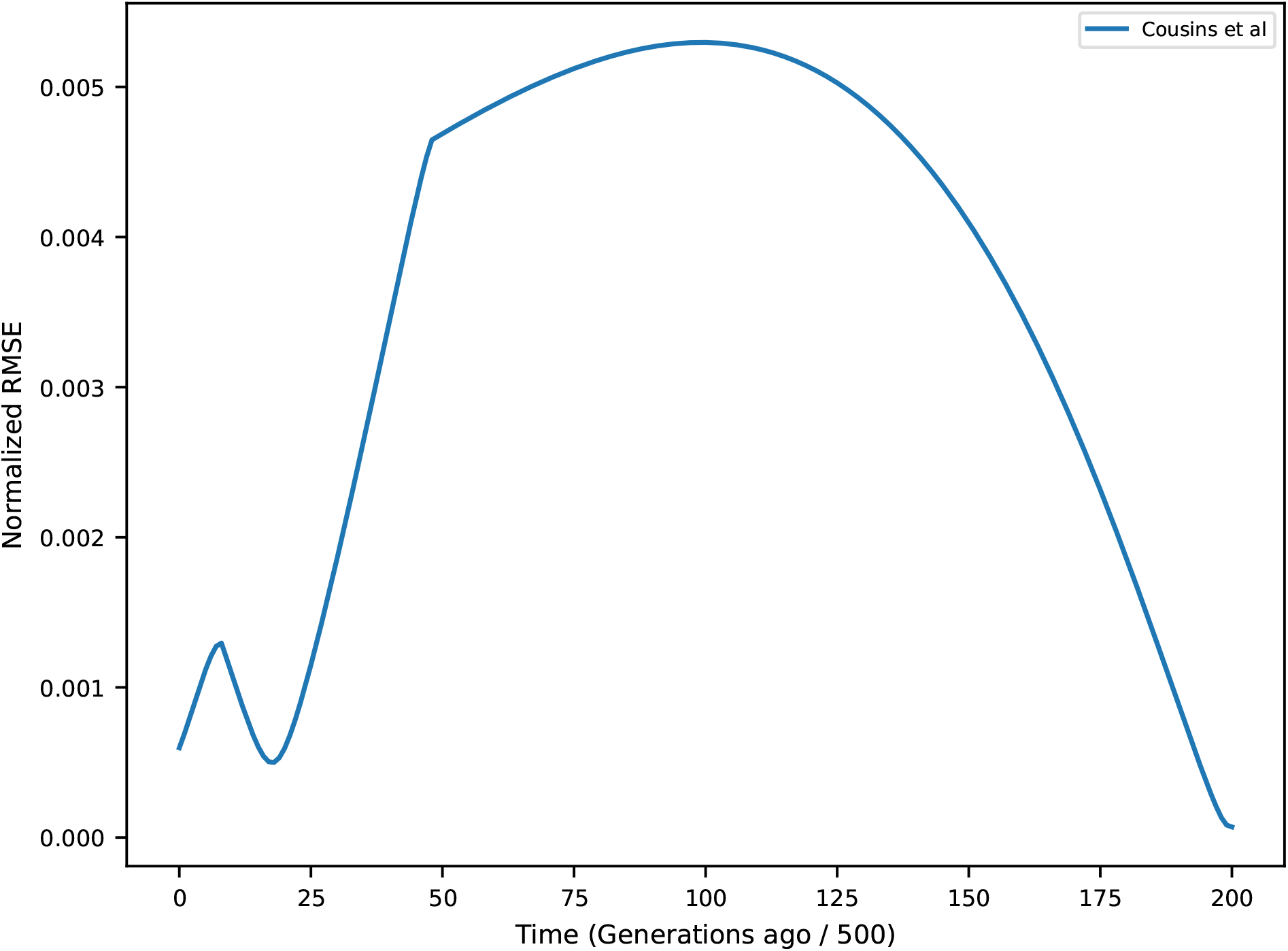
Temporal evolution of the the Normalized Rooted Mean Squared Error between the demography-aware *B*-map and steady-state *B*-maps, predicted every 500 generations after the first size change in the Cousins et al. (2024) demography. Steady-state *B*-maps are obtained using the equilibrium *N*_*e*_ value calculated from neutral genetic diversity in the absence of linked selection (equivalent to the harmonic mean of population sizes in the relevant time-frame). Here the chromosome layout follows that of Figure 3.

**Figure S6:**
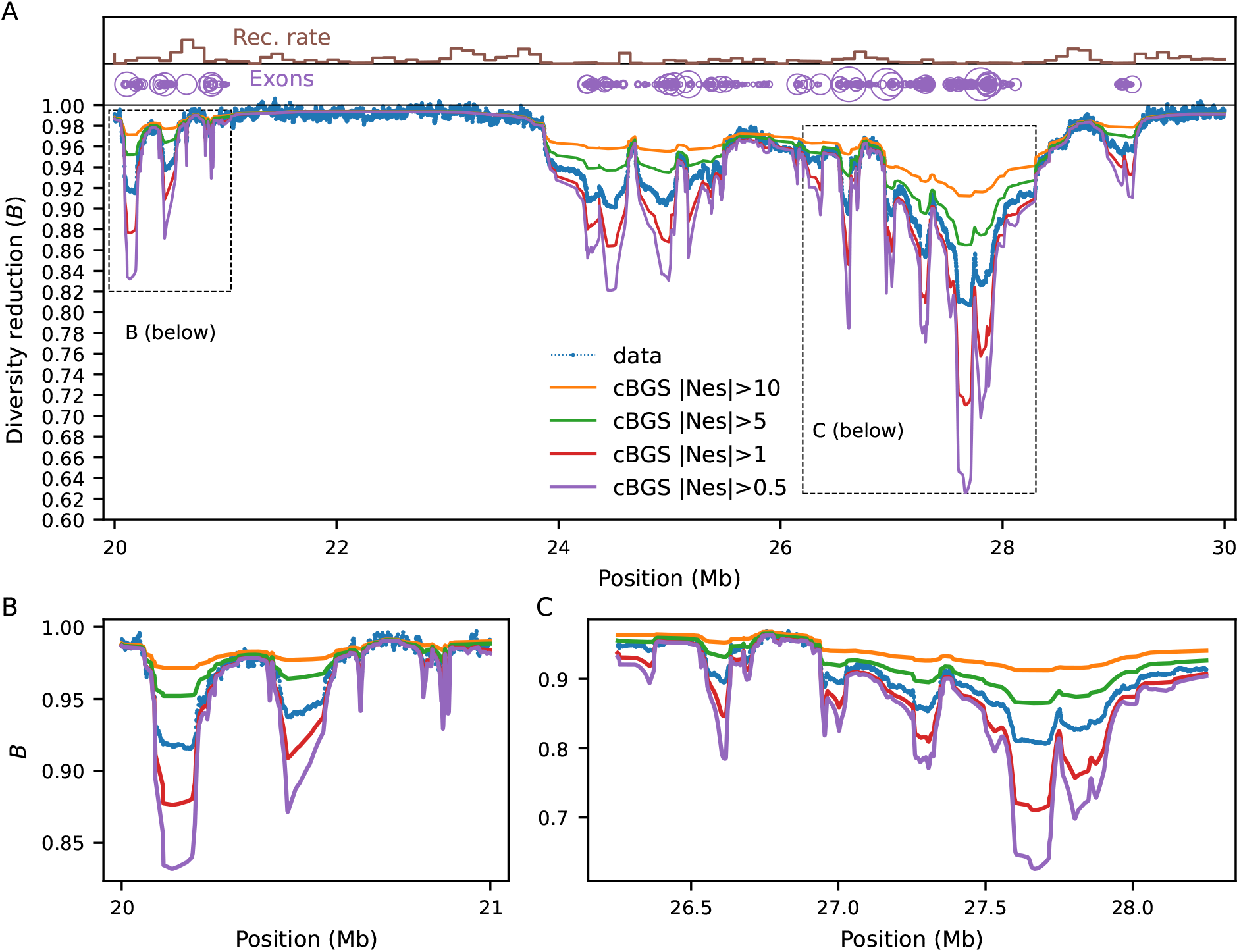
Predictions from cBGS, truncating the DFE at different points. Notice the overestimation of *B*-values caused by excluding a large portion of the DFE (orange and green lines) as well as the underestimation caused by the inappropriateness of the cBGS model at weaker selection coefficients (red and purple lines). Here the chromosome layout follows that of Figure 3.

**Figure S7:**
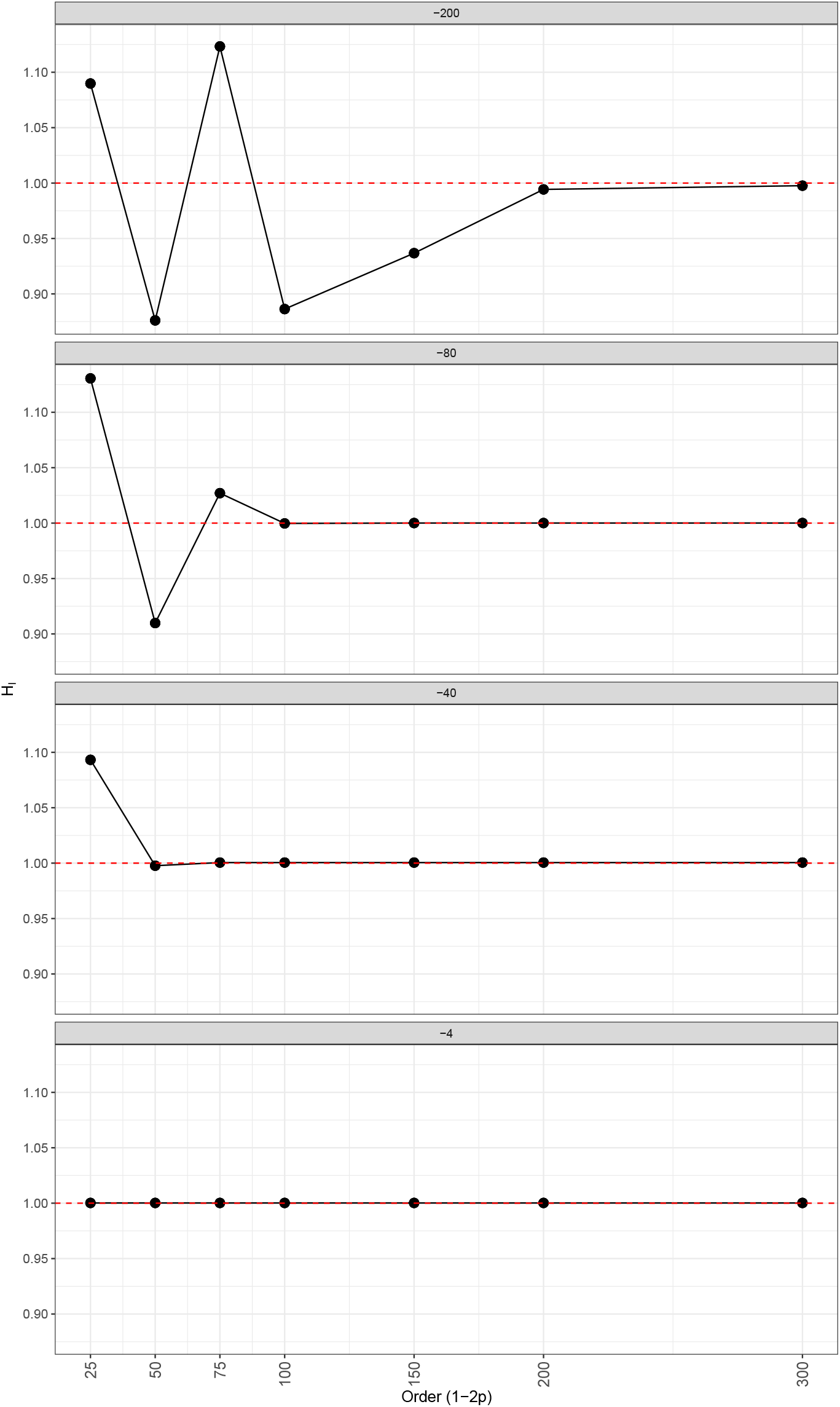
Benchmarking of the implementation of the selection operator in moments++, with the truncation strategy to close the system. Shown are ratios of *π*_*l*_ predicted with moments++ to *π*_*l*_ predicted with moments.TwoLocus, as a function of the Order of 1 − 2*p* factors included in the moments++ model (panels show *N*_*e*_*s*). Dashed red lines denote the target line indicating good agreement with our gold-standard.

**Figure S8:**
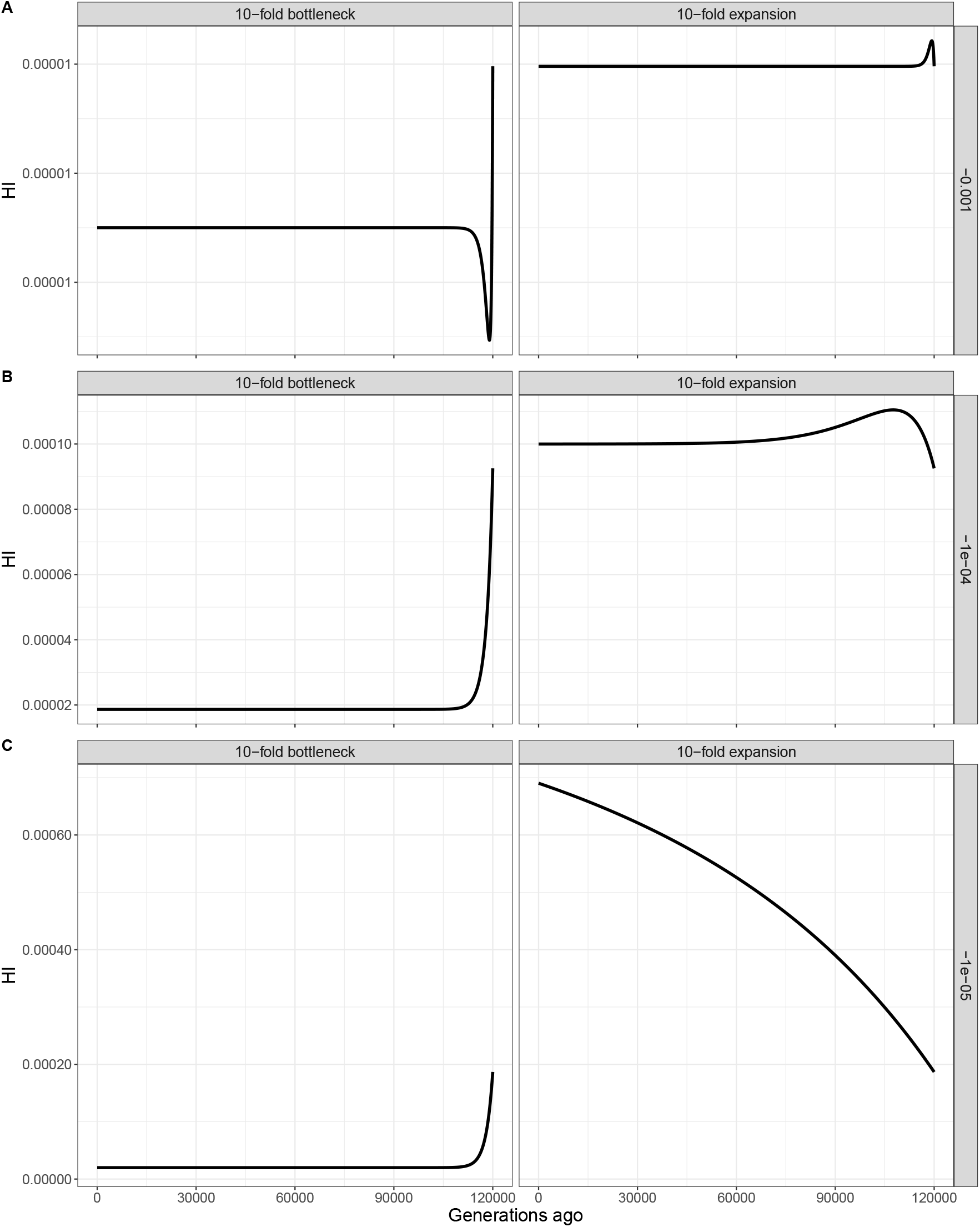
Temporal trajectories of *π*_*L*_ following a 10-fold bottleneck (left panels) or expansion (right panels). A) *s* = −0.001. B) *s* = −0.0001. C) *s* = −0.00001. Here *µ* = 10^−8^.

**Figure S9:**
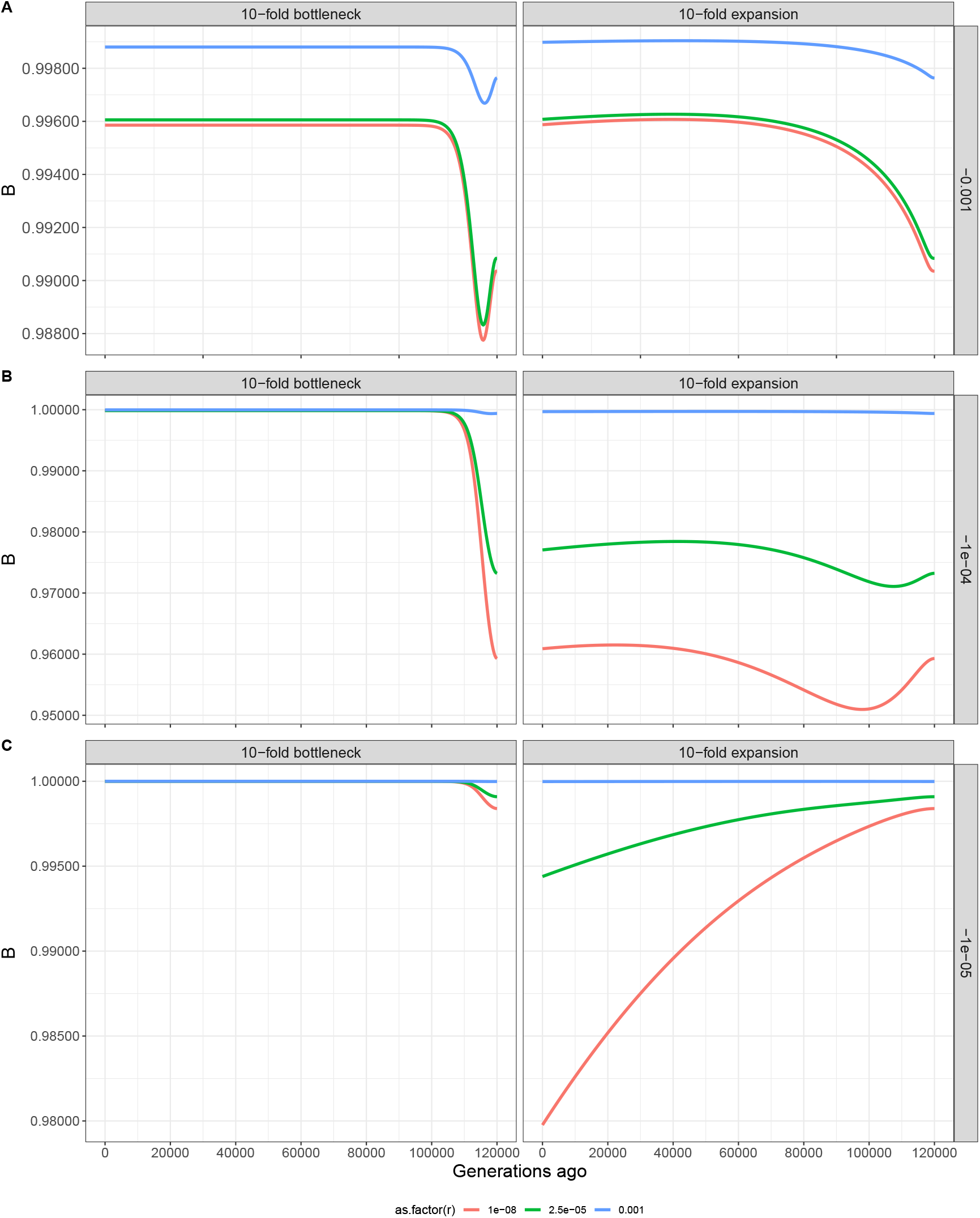
Temporal trajectories of *B*-values following a 10-fold bottleneck (left panels) or expansion (right panels), for a pure two-locus model. Note how the expansion scenario does not reach the new equilibrium after 120,000 generations. A) *s* = −0.001. B) *s* = −0.0001. C) *s* = −0.00001. Colors denote the rate of recombination. Here *µ* = 10^−8^ and *B*-values are raised to the 1000th power to facilitate visualization.

**Figure S10:**
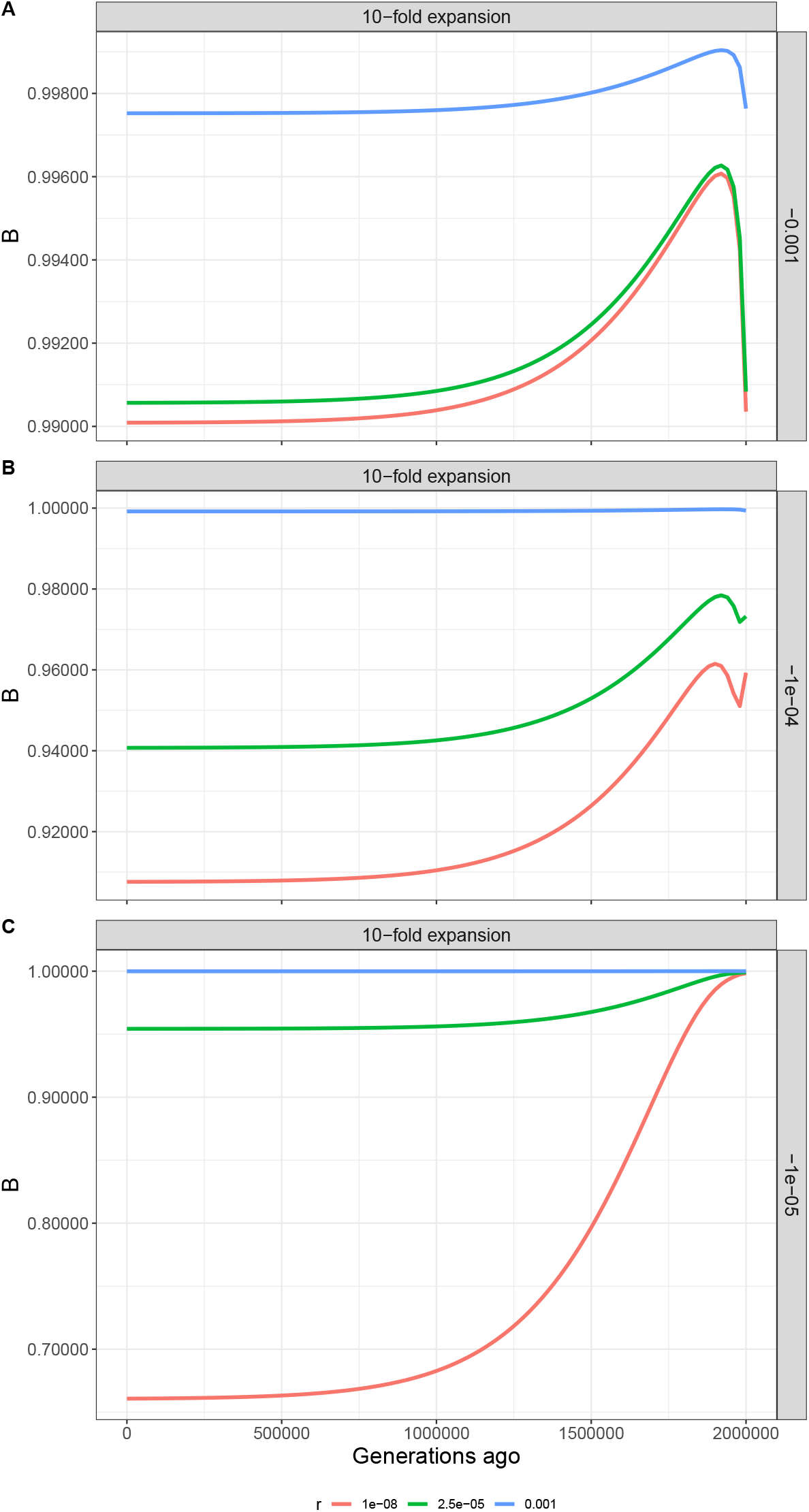
Temporal trajectories of *B*-values under the 10-fold expansion model from Figure S9 where we track statistics for 2,000,000 generations to ensure that the new equilibrium point is found. A) *s* = −0.001. B) *s* = −0.0001. C) *s* = −0.00001. Colors denote the rate of recombination. Here *µ* = 10^−8^ and *B*-values are raised to the 1000th power to facilitate visualization.

**Figure S11:**
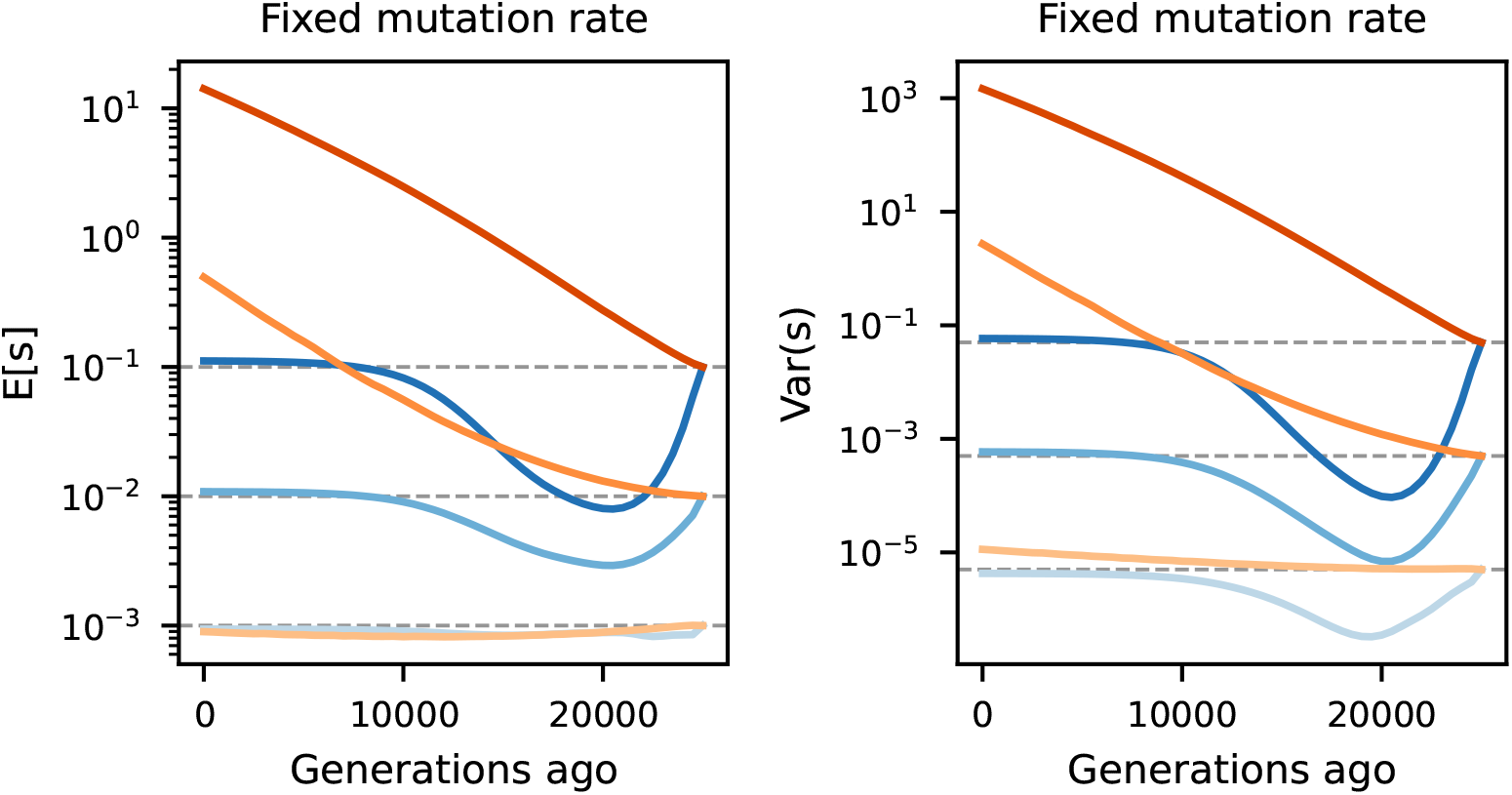
Mean (left) and variance (right) of the inferred DFEs depicted in Figure 4.

**Figure S12:**
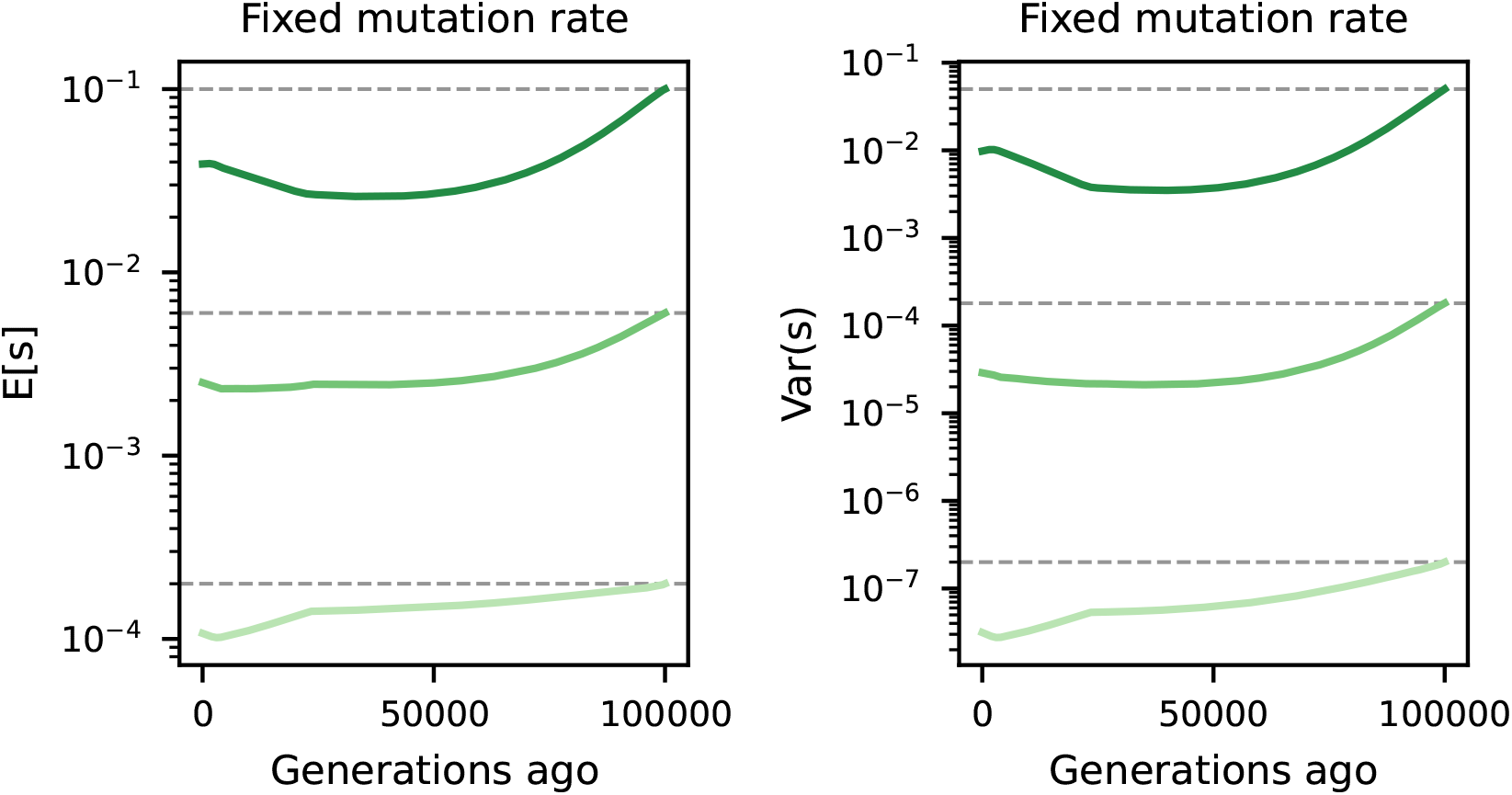
Mean (left) and variance (right) of the inferred DFEs depicted in Figure 5.

